# Development of a multi-primer metabarcoding approach to understand trophic interactions in agroecosystems

**DOI:** 10.1101/2021.05.19.444782

**Authors:** Ivan Batuecas, Oscar Alomar, Cristina Castañe, Josep Piñol, Stéphane Boyer, Lorena Gallardo-Montoya, Nuria Agustí

**Author notes:** Correspondence to: Iván Batuecas, IRTA, Ctra. de Cabrils, Km. 2, E-08348 Cabrils, Barcelona, Spain.

## Abstract

Knowing which arthropod and plant resources are used by generalist predators in agroecosystems is important to understand trophic interactions and the precise ecological role of each predatory species. To achieve this objective, molecular approaches, such as the use of high-throughput sequencing (HTS) platforms are key. This study develops a multi-primer metabarcoding approach and explores its suitability for the screening of the most common trophic interactions of two predatory species of arthropod with contrasted morphology, *Rhagonycha fulva* (Coleoptera: Cantharidae) and *Anthocoris nemoralis* (Hemiptera: Anthocoridae) collected in an organic peach crop. To save time and cost in this metabarcoding approach, we first evaluated the effect of two different predator-pool sizes (10 and 23 individuals of the same species), as well as the performance of using one or two primer pairs in the same library. Our results show that the analysis of 23 individuals together with the use of two primer pairs in the same library optimizes the HTS analysis. With these best-performing conditions, we analyzed whole bodies of field-collected predators as well as the washing solutions used to clean the insect bodies. Results showed that we were able to identify both, gut content (i.e. diet) as well as external pollen load (i.e. on the insects’ body), respectively. This study also demonstrates the need of washing predatory insects prior to HTS analysis when the target species have a considerable size and hairy structures. This metabarcoding approach has a high potential for the study of trophic links in agriculture, revealing both expected and unexpected trophic relationships.

## Introduction

The management of ecosystem services in agroecosystems is key for food production. One of these ecosystem services is pest control, carried out by natural enemies, such as insect generalist predators. Commonly, these beneficial insects do not only require prey as food, but they also need plant resources as food and/or as habitat supply (Demestihas et al. 2017). A more thorough understanding of how generalist insect predators use these resources in an agroecosystem is important to further utilize these predators in pest control programs.

Studying trophic interactions within an ecosystem is inherently difficult, because predation is an ephemeral process often difficult to visualize. Different methods have been used to measure insect predation, from their direct observation in the field, to the molecular analyses of their gut contents (Agustí et al. 2003; Pumariño et al. 2011; Nielsen et al. 2018). Molecular approaches to study predation increases the precision of the diet description (Nielsen et al. 2018), particularly with the use of high-throughput sequencing (HTS) platforms, which allow the detection of more realistic trophic interactions conducted in the field. Within these HTS (also called next generation sequencing or NGS) approaches, DNA metabarcoding, understood as the identification of organisms from a sample containing DNA from more than one organism, has been used to describe interactions in both terrestrial and aquatic ecosystems (Kennedy et al. 2020). Metabarcoding can be very helpful in agroecosystems, particularly for an initial screening of the gut content analysis of generalist predators (Pompanon et al. 2012), as already shown in few other cases (Piñol et al. 2014; Gomez-Polo et al. 2015, 2016; González-Chang et al. 2016).

DNA metabarcoding studies usually follow a well-established workflow that includes the DNA extraction often from the whole specimens, a PCR amplification with barcoded primers, high-throughput DNA sequencing, and a tailored bioinformatic analysis to obtain the desired taxonomic classification (Deagle et al. 2018). Nevertheless, recent literature highlights that several factors can affect the final result, indicating that certain technical aspects need to be improved (Lamb et al. 2019). These factors include the need for an external washing of the predator specimens to remove foreign external contamination (e.g. pollen grains) from their exoskeleton (Jones, 2012); the need for pooling samples, particularly when ingested DNA template is low; the use of biological replicates to obtain robust estimates of diet diversity and composition (Mata et al. 2019); the number of primer pairs used (Gibson et al. 2014); the availability of comprehensive reference databases with regards to the taxonomic groups of interest (Bohmann et al. 2011); or the use of different pipelines and data cleaning procedures during the bioinformatic analysis (Plummer et al. 2017). The use of more than one primer set has been previously recommended in order to minimize the effect of set biases and to recover a higher taxonomic coverage of the diet (Piñol et al. 2015; Krehenwinkel et al. 2017; Hajibabaei et al. 2019). With that in mind, we developed a new metabarcoding approach using two arthropod and two plant universal primer pairs per library to describe the main consumed taxa of predator diets by HTS, and we have tested it on two generalist insect predator species.

The main aim of this study was to explore the suitability of a multi-primer metabarcoding approach to provide a screening of the most common trophic interactions in the agroecosystem with pooled samples, whilst considering the reduction on time and cost when field-collected predatory arthropod specimens have to be analysed. We focused on two predator species, the minute pirate bug *Anthocoris nemoralis* (Fabricius) (Hemiptera: Anthocoridae), and the common red soldier beetle *Rhagonycha fulva* (Scopoli) (Coleoptera: Cantharidae). Both insects are present in peach crops in Lleida region (NE Spain), as well as in other fruit and arable crops in the same area of study, like maize or alfalfa (Pons & Eizaguirre, 2000; Jauset et al. 2007). *Anthocoris nemoralis* is known as one of the most important biocontrol agents of the pear psyllids *Cacopsylla pyricola* (Foerster) and *Cacopsylla pyri* L. (Hemiptera: Psyllidae) (Agustí et al. 2003). However, this predatory species has also been described to feed on pollen (Naranjo & Gibson 1996). *Rhagonycha fulva* is mainly present in wooded agricultural landscapes and arable lands (Meek et al. 2002; Rodwell et al. 2018). Even if this species is mainly known to feed on pollen and nectar from umbellifers (Apiaceae) (Meek et al. 2002), it has also been cited as predator of some insect species (Pons & Eizaguirre, 2000; Rodwell et al. 2018). Nevertheless, its role as biocontrol agent is not well-known, as it is also the case for *A. nemoralis* in other fruit crops than pears, like peaches. The selected predator species are morphologically different regarding their potential to retain pollen grains on their body. *Rhagonycha fulva* is large (10-15 mm) and pubescent, particularly on its head and ventral side, while *A. nemoralis* is much smaller (3 mm) and glabrous. These different morphological characteristics make them good candidates to study pollen retention on their bodies, and therefore the need of washing them before HTS analysis.

In this study, we have investigated the effect of a variable sample-pool size on the range of prey taxa detected (taxonomic coverage); as well as the effect of using one or two primer pairs in the same library. We then validated the developed methodology by analysing the arthropod and plant diet of two small populations of *A. nemoralis* and *R. fulva*, two omnivorous insects with contrasted morphology. Plant and other arthropod DNA content in their washing solutions was also analyzed as a mean to identify the pollen present on their body while foraging on diverse plants in the landscape.

## Materials and Methods

### Predator collection, cleaning and DNA extraction

*Anthocoris nemoralis* (n=42) and *R. fulva* (n=78) were collected by beating branches in a peach orchard in Vilanova de Segrià (Lleida), Spain (UTM 10×10: 31TCGO1) in June and July 2016, and May 2017, respectively. Each specimen was individualized in a DNA-free tube and placed in a portable freezer to avoid DNA degradation. Once in the lab, specimens were morphologically identified and stored at -20°C until metabarcoding analysis.

Before DNA extraction, all collected specimens were individually washed in order to remove contaminants from their cuticle. The washing process consisted in submerging each insect in a 10 ml tube containing a DNA-free water solution with Tween^®^ 20 (0.1%) and manually shaking the tube for 1 min. This washing solution was stored at -20°C for further HTS analysis (see below the *Analysis of field-collected predators* section). After that, the insect was submerged in another 10 ml tube with DNA-free water solution containing sodium hypochlorite (0.5 %) and Tween^®^ 20 (1%) and the tube shaken manually for another 1 min. This second washing solution was discarded. Finally, each insect was rinsed with DNA-free water for 30 seconds and dried on filter paper.

The DNA of each insect specimen or washing solution was extracted using the Speedtools Tissue DNA Extraction Kit (Biotools, Germany; protocol for animal tissues). DNA from washing solutions was extracted with an additional disruption step using 0.15 g of 500–750 μm diameter glass beads (ACROS Organics™), and vortexed for 15 min at 50 Hz in a Gene2 vortex (MoBio Laboratories), for a suitable breakage of the potentially present pollen grains. Plastic pestles were used for whole insects instead. After the DNA extraction process, total DNA was eluted in 100 µl of AE buffer provided by the manufacturer and stored at −20°C. A negative control without insect or plant DNA (DNA-free water) was added to each DNA extraction set. The concentration of each DNA extraction was measured on a Qubit^®^ 2.0 fluorometer using the dsDNA HS Assay kit (Invitrogen, Carlsbad, CA, USA). Equimolar amounts of each individual DNA extraction (5 ng/µl) were finally pooled by species in sample-pools, as shown in Table 1.

**Table 1.**
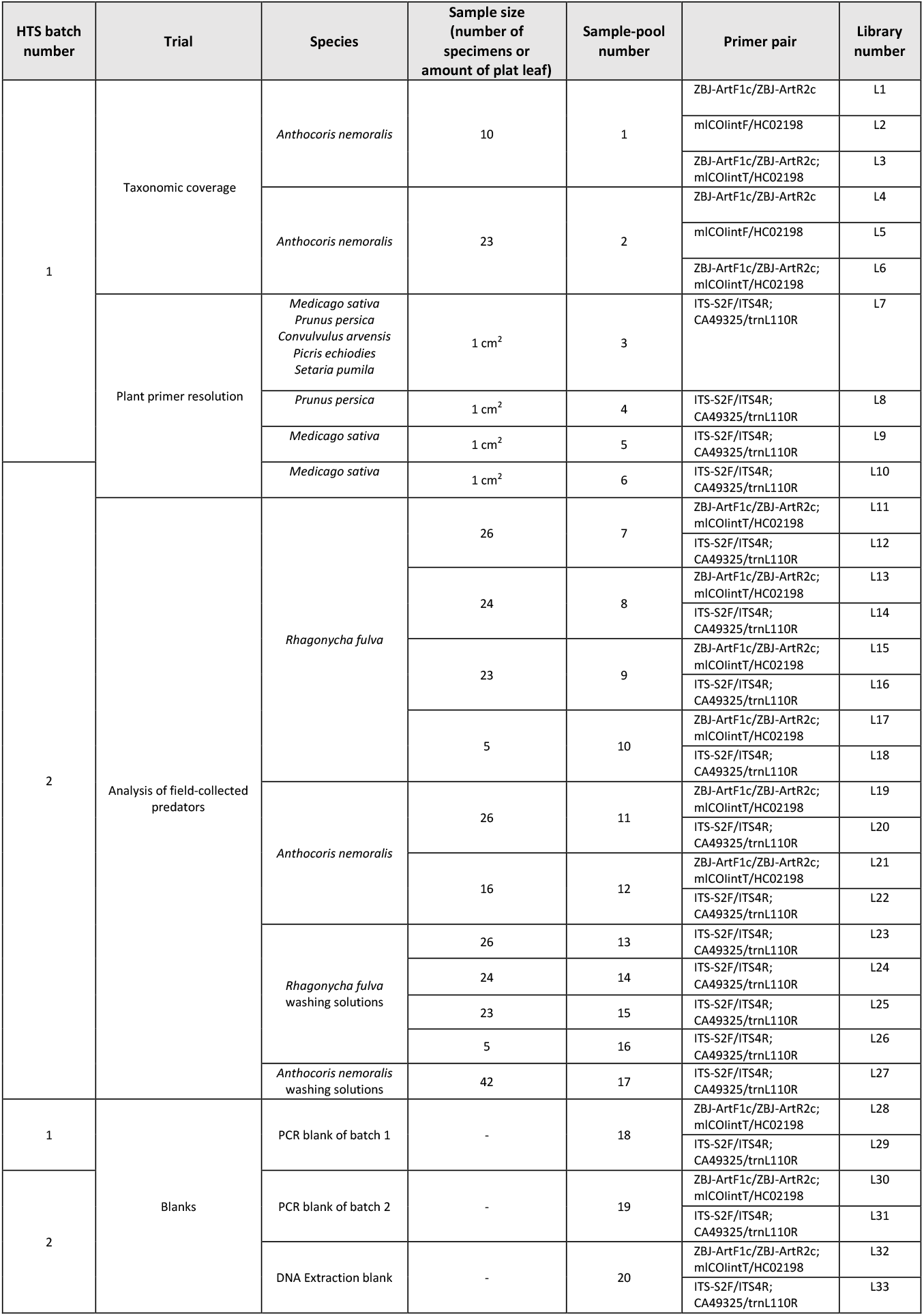
Arthropod and plant sample-pools analysed by HTS in the three trials conducted in the present study. Also indicated the number of the HTS batch where each library was analysed, the number of individuals (or amount of plant material) in each sample-pool, and the primer pairs used in each library. ZBJ-ArtF1c/ZBJ-ArtR2c and mlCOIintF/HC02198 are the arthropod universal primers used, and ITS-S2F/ITS4R and CA49325/trnL110R are the plant universal primers used.

### PCR amplification, library preparation and sequencing

Two pairs of universal arthropod primers which partially amplify the mitochondrial Cytochrome Oxidase subunit I (COI) region were used to amplify DNA from the field-collected insects. These two pairs of primers were selected because they amplify different amplicon sizes (ZBJ-ArtF1c/ZBJ-ArtR2c, 157 bp; and mlCOIintF/HC02198, 313 bp) and do not overlap in the COI region (Table S1; Fig. S1), thereby avoiding competition for the same primer binding sites. Similarly, we used two pairs of universal plant primers also amplifying different amplicon sizes (ITS-S2F/ITS4R, 350 bp; and cA49325/trnL110R, 80 bp) (Table S1). In this case, primer pairs were chosen to amplify fragments in different regions, the first in the nuclear Internal Transcribed Spacer 2 (ITS2) and the second in the chloroplast *trnL* intron.

Sample-pools shown in Table 1 were amplified using a universal multi-primer approach with these four pairs of universal primers for arthropods and plants, performing one PCR with both pairs of arthropod primers, and another one with both pairs of plant primers. Each PCR reaction (50 µL) contained 25 µL of Multiplex Master Mix (Qiagen, Hilden, Germany), 1 µL of each primer [10 μM], 8 µL of free-DNA water and 15 µL of DNA of each sample-pool. PCR conditions used with the arthropod primers were: 95 °C for 5 min for the initial denaturation, followed by 30 cycles at 95 °C for 30 s, 46 °C for 30 s and 72 °C for 30 s, and a final extension at 72 °C for 10 min. PCR conditions used with the plant primers were: 95 °C for 3 min, followed by 30 cycles at 95 °C for 30 s, 55 °C for 30 s and at 72 °C for 30 s, and a final extension at 72 °C for 5 min. Amplifications were conducted in a 2720 thermocycler (Applied Biosystems, CA, USA). Target DNA and DNA-free water were included in each PCR run as positive and negative controls, respectively. Resulting PCR products were purified with QIAquick PCR Purification kit (Qiagen), and 5 µl of each PCR product was used afterwards as template to prepare the libraries to be sequenced. HTS analysis was conducted in two batches (Table 1), and libraries of both batches were built by mixing the PCR products of either both pairs of arthropod primers, or both pairs of plant primers. Both HTS batches were processed on a MiSeq sequencing platform (Illumina, San Diego, CA, USA) at the *Servei de Genòmica i Bioinformàtica* of the Autonomous University of Barcelona, Spain. Illumina adapters were attached using Nextera XT Index kit. Amplicons were purified with magnetic beads and 5 µl of each library were pooled and sequenced with a paired-end approach (2 × 225 bp).

### Taxonomic coverage: sample-pool size and number of arthropod primer pairs

Two different sample-pools of *A. nemoralis* were build: sample-pool 1, with 10 individuals; and sample-pool 2, with 23 individuals (Table 1, *Taxonomic coverage*). Only in this trial, both sample-pools were tested using either both universal arthropod primer pairs together in the same library (L3 and L6) or separated in different libraries (L1, L2, L4 and L5). The effects of the sample-pool size (sample-pool 1 vs 2) and the use of one or both primer pairs together in the same library on the number of taxa obtained (taxonomic coverage) after HTS was compared using the non-parametric Kruskal-Wallis rank sum test. The statistical analyses were performed with R version 3.5.1 (R Development Core Team, 2018).

### Plant primer resolution

To test the efficacy of each pair of plant primers and to assess their level of taxonomic resolution, we built a plant sample-pool with five plant species that are common in orchard ground covers and field margins of the study area (Table 1, sample-pool 3) (Ibáñez-Gastón, 2018; Juarez-Escario et al. 2010). In addition, to validate the accurate parameterization of the bioinformatic pipeline (Jusino et al. 2019), we included three positive controls containing only the crop plants (sample-pools 4-6). Unlike arthropods, plant samples (1 cm^2^ leaf disc) were not washed prior to DNA extraction, which was conducted in the same way as for arthropod samples.

### Analysis of field-collected predators

Field-collected predators were analysed with the multi-primer approach described above and using the most appropriate sample-pool size and number of primer pairs, according to the results of the previous *Taxonomic coverage* and *Plant primer resolution* trials. Four sample-pools were tested for *R. fulva* (Table 1, sample-pools 7-10) and two for *A. nemoralis* (sample-pools 11 and 12). In addition, five sample-pools were analysed in order to identify pollen load on the insects’ body: four sample-pools from *R. fulva* washing solutions (sample-pools 13-16), and one from *A. nemoralis* washing solutions (sample-pool 17). In order to determine whether both predators only foraged on plants or also consumed plant resources, we compared the obtained plant taxa from their washing solutions with those obtained from their gut contents. Finally, in order to increase the amount of taxa detetected with the aim to show the highest diet diversity, we have considered each sample-pool of the same predator species as a different biological replicate, which provides greater variability than technical replicate for the taxa detected (Mata et al. 2019).

### Bioinformatics

Raw Illumina reads were merged using VSEARCH 2.0 algorithm (Rognes et al. 2016), and then analysed using a restrictive strategy to reduce biases. The assembled reads were quality filtered using the FASTX-Toolkit tool (Gordon & Hannon, 2010) with a minimum of 75% of bases ≥Q30. The resulting reads were then split by length according to the expected amplicon from each primer with custom Python scripts. Primer sequences were removed using Cutadapt 1.11 (Martin, 2017). The obtained reads were clustered into OTUs with a similarity threshold of 97% using VSEARCH 2.0. Chimeras were removed using the UCHIME algorithm (Edgar et al. 2011). The remaining OTUs were queried against custom-made databases using BLAST 2.2.31+ (BLASTN, E-value 1e-10, minimum coverage of the query sequence: 97%, numbers of alignments: 9) (Camacho et al. 2009). The custom-made databases contained all arthropod and plant sequences present in the study area and available in the NCBI database (http://www.ncbi.nlm.nih.gov) at the moment of the analysis (October 2019). For this, we used European and regional biodiversity databases: GBIF.org (http://www.gbif.org/) and *Banc de dades de biodiversitat de Catalunya* (http://biodiver.bio.ub.es/biocat/). Taxonomy was assigned at ≥97% identity by Last Common Ancestor algorithm (LCA) with BASTA (Kahlke & Ralph 2019). To remove possible contaminants from the OTUs assigned to different taxa for each group of primer pairs (arthropods or plants), we considered in the analysis only those OTUs that strictly had more than five reads and that had been detected in at least two sample-pools of the same species (Boyer et al. 2013). When the OTUs were obtained only in one sample-pool, they were used in the analysis only if they had more than five reads with both primer pairs, or if they exceeded the 0.03 % threshold of the total reads for plant or arthropod in each case. Obtained OTUs were then categorized as predator or prey based on their taxonomy.

To reduce other biases, such as secondary predation (an important limitation of HTS when studying food webs (da Silva et al. 2019)), and also with the aim of showing the most important taxa ingested, dietary data were presented using two dietary metrics, as recommended by Deagle et al. (2018). The first metric was the percentage of Relative Read Abundance (RRA), which was calculated considering the total number of reads of each consumed resource (arthropod or plant) amplified with each primer pair and for each library, divided by the number of total reads of all resources obtained with each primer pair for each library. After that, a filter to eliminate resources <1% of the amplified taxa was applied, as recommended by Deagle et al. (2018). This was applied for each primer pair in each library. With the taxa obtained, the second metric was calculated, which was the percentage of Frequency of Occurrence (FOO), being the percentage of the total number of pools of each specimens analysed that contain a resource items obtained, indicating the most common resources consumed.

## Results

The analysis of 33 libraries (120 predators and 20 sample-pools) conducted in two HTS batches (Table 1) generated 9,047,294 raw paired-end reads, 95% of which were successfully merged (Table 2, step 1). After that, 40,582 (step 2) and 53,286 reads (step 3) that did not match our quality and/or length requirements were discarded, as well as 2,512 chimera reads (step 5). After the taxonomic assignment (step 6), 1,548 arthropod and 649 plant OTUs were filtered (step 7 and Step 8). From the initial raw paired-end reads, only 8,930 (0.098%) came from the DNA extraction blank (sample-pool 20) and both PCR blanks (sample-pools 18 (batch 1) and sample-pool 19 (batch 2)) (Table 1). Those reads were eliminated at the step 7. After calculating RRA and FFO percentages and eliminating taxa with a number of reads lower than 1% (Table 2, step 8; Table S2), we finally obtained 299 arthropod and 206 plant OTUs (Table 2), which corresponded to 14 arthropod and 20 plant taxa (Table 3).

**Table 2.**
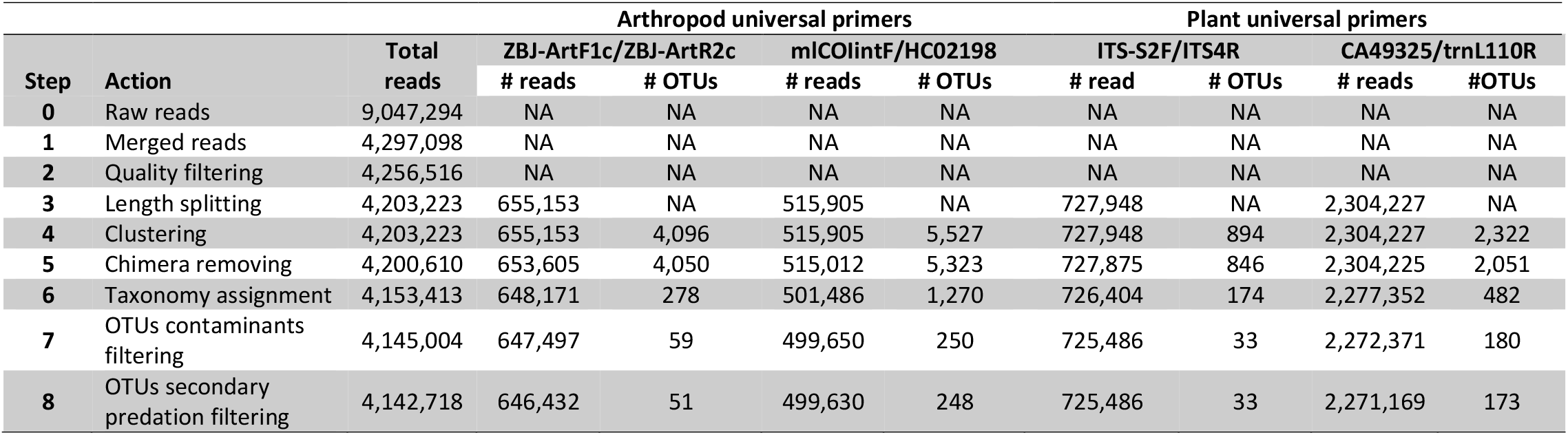
Number of reads and OTUs obtained at each step of the bioinformatic analysis. Data is presented in total and according to each arthropod and plant primer pair in each step of the bioinformatic analysis. NA = not applicable.

**Table 3.** Summary table of all arthropod (n=14) and plant (n=20) taxa obtained after bioinformatic analysis of HTS data (33 libraries of 20 different sample-pools (see Table 1)). The lowest taxonomic rank reached is indicated in bold.

| Kingdom | Phylum | Order | Family/Subfamily | Genus | Species |
| --- | --- | --- | --- | --- | --- |
| Animalia | Arthropoda | Hemiptera | Anthocoridae |  | *Anthocoris nemoralis* Fabricius |
|  |  |  |  | *Orius* |  |
|  |  |  |  |  | *Orius laevigatus* Fieber |
|  |  |  | Aphididae |  |  |
|  |  |  |  |  | *Myzus persicae* Sulzer |
|  |  |  | Lygaeidae |  | *Nysius graminicola* Kolenati |
|  |  |  | Liviidae |  | *Diaphorina lycii* Loginova |
|  |  | Coleoptera | Coccinellidae |  |  |
|  |  |  |  |  | *Oenopia conglobata* L. |
|  |  |  | Cantharidae |  | *Cantharis livida* L. |
|  |  |  |  |  | *Rhagonycha fulva* Scopoli |
|  |  | Diptera | Cecidomyiinae |  |  |
|  |  | Lepidoptera | Tortricidae |  | *Grapholita molesta* Busck |
|  |  | Thysanoptera | Thripidae |  | *Thrips fuscipennis* Haliday |
| Plantae | Streptophyta |  |  |  |  |
|  |  | Asterales | Asteraceae |  |  |
|  |  |  |  | *Sonchus* |  |
|  |  |  |  |  | *Picris echioides* L. |
|  |  | Solanales | Convolvulaceae |  |  |
|  |  |  |  |  | *Convolvulus arvensis* L. |
|  |  | Solanales | Solanaceae |  |  |
|  |  | Fabales | Fabaceae |  |  |
|  |  |  |  |  | *Medicago sativa* L. |
|  |  |  |  | *Trifolium* |  |
|  |  | Lamiales | Oleaceae |  | *Olea europaea* L. |
|  |  | Pinales | Pinaceae | *Pinus* |  |
|  |  | Poales | Poaceae |  |  |
|  |  |  |  | *Setaria* |  |
|  |  |  |  |  | *Dactylis glomerata* L. |
|  |  |  |  |  | *Poa annua* L. |
|  |  | Caryophyllales |  |  |  |
|  |  |  | Amaranthaceae |  | *Beta vulgaris* L. |
|  |  | Rosales | Rosaceae |  |  |
|  |  |  |  |  | *Prunus persica* (L.) Batsch |

### Taxonomic coverage: sample-pool size and number of arthropod primer pairs

The six libraries analysed in this trial (Table 1, L1-L6), yielded 10 arthropod taxa (Table S3; Table 3). Besides the predator itself (*A. nemoralis*), we detected other anthocorids (*Orius* and *O. laevigatus* Fieber), other potential predator (Cecidomyiinae), as well as some pest (Aphididae, *Grapholita molesta* Busck, *Myzus persicae* Sulzer (Aphididae), *Thrips fuscipennis* Haliday) and non-pest prey (*Diaphorina lycii* Loginova).

The number of arthropod taxa obtained was not significantly different between libraries mode of 23 or 10 *A. nemoralis* individuals (Kruskal-Wallis chi-squared = 0.78, df = 1, p-value = 0.37) (Table S4). Similarly, when comparing the number of arthropod taxa obtained using only one or two pairs of primers together in the same library, no significant differences were observed (Kruskal-Wallis chi-squared = 0.16, df = 1, p-value = 0.68) (Table S4). Therefore, in order to save time and cost in the following *Analysis of field-collected predators* trial, we decided to pool up to 26 predators together, and to use both pairs of arthropod primers together in the same library.

### Plant primer resolution

The four plant libraries analysed in this trial (Table 1, L7-L10), yielded 11 plant taxa (Table S3; Table 3). Most of these taxa were expected because they were present in the composition of the sample-pools 3-6 (Table 1) (*Medicago sativa* L. (alfalfa), *Prunus persica* (L.) (peach), *Convulvulus arvensis* L., *Picris echioides* L., *Setaria* sp.), which were used as positive controls. Other plant taxa were also detected, like Streptophyta, which corresponds to a clade that show just plant DNA amplificated without additional taxonomic level information, and the families Fabaceae, Rosaceae, Convolvulaceae and Asteraceae, which were the families of the plant species of the sample-pools 3-6 (Table S3; Table 3). The genus *Trifolium* (Fabaceae) in the library L7 was also detected (Table S3). Nevertheless, it represented only 0.026% of the total reads obtained, and for this reason, it was not considered in further analysis.

### Analysis of field-collected predators

The 17 libraries analysed in this trial (Table 1, L11-L27), yielded 28 taxa (14 of arthropods and 14 of plants (Table S3; Fig. 1)). Regarding the diet of *R. fulva* (Cantharidae), besides the predator itself, we detected three other arthropod taxa: *Nysius graminicola* Kolenati (Lygaeidae), *Cantharis livida* L. (Cantharidae) and Coccinellidae; and five plant taxa: Streptophyta, Convolvulaceae, Solanaceae, Fabaceae and Poaceae (Table S3; Fig. 1; Table 3). Regarding the diet of *A. nemoralis* (Anthocoridae), besides the predator itself, we detected 9 other arthropod taxa: *Orius* and *O. laevigatus* (Anthocoridae), Aphididae, *M. persicae* (Aphididae), *D. lycii* (Liviidae), *Oenopia conglobata* L. (Coccinellidae), Cecidomyiinae, *G. molesta* (Tortricidae) and *T. fuscipennis* (Thripidae). No plant taxa were obtained in this HTS analysis from whole specimens of *A. nemoralis* (Table S3; Fig. 1).

**Figure 1.**
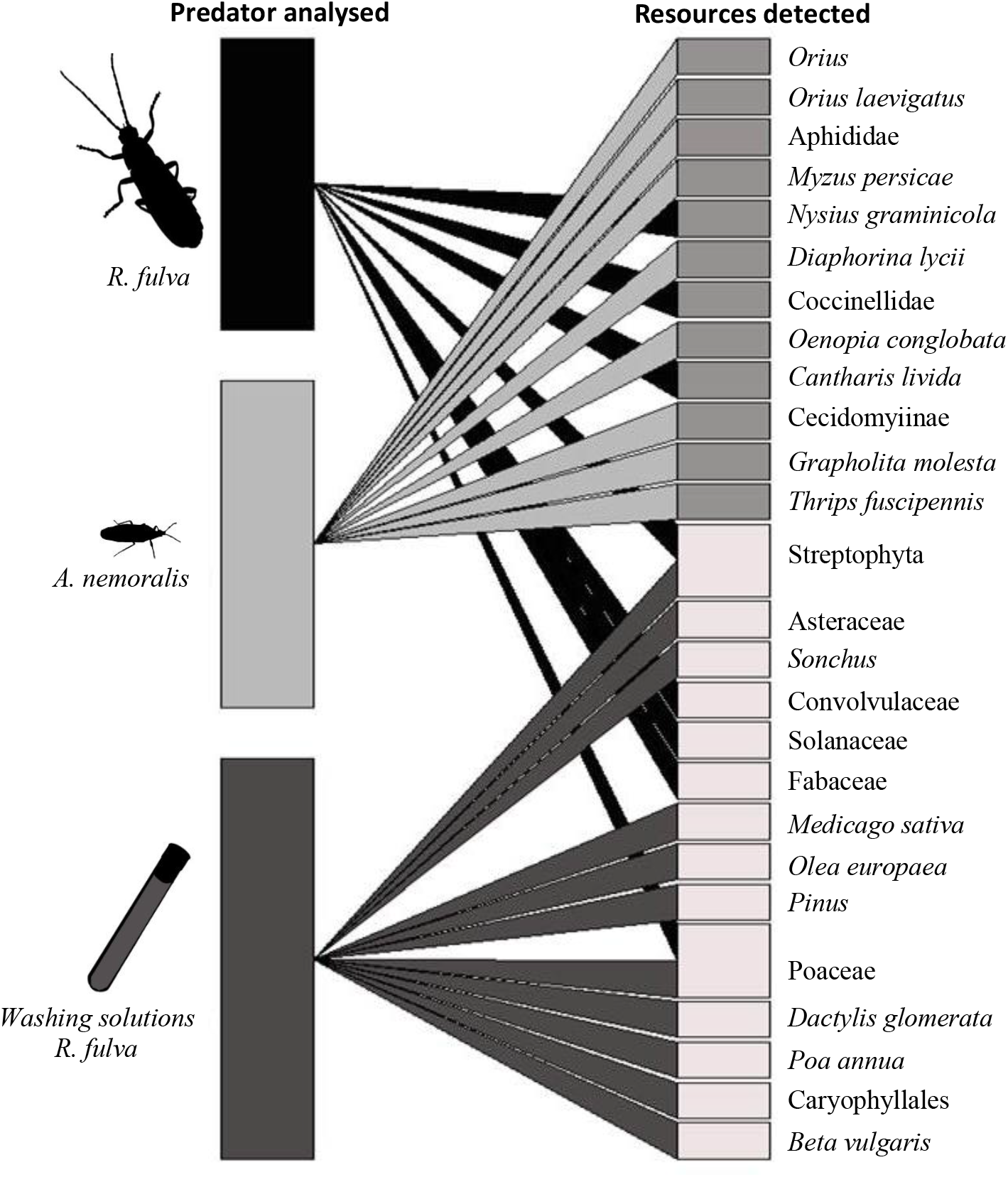
Interaction network of the arthropod and plant taxa detected from whole body extractions of *Ragonycha fulva* and *Anthocoris nemoralis*, as well as from the washing solutions of *R. fulva*.

We obtained amplification in two of the four libraries from the washing solutions of *R. fulva* analysed (Table 1, L23-L26) (Table S3). The 11 plant taxa detected were: Streptophyta, Asteraceae, *Sonchus* (Asteraceae), *M. sativa* (Fabaceae), *Olea europea* L. (Oleaceae), *Pinus sp* (Pinaceae), Poaceae, and *Dactylis glomerata* L. and *Poa annua* L. (Poaceae), Caryophyllales, *Beta vulgaris* L. (Amaranthaceae) (Table S3; Fig. 1). No plant taxa were detected from *A. nemoralis* washing solutions (Table 1, L27; Table S3).

## Discussion

### Methodological issues

The present study addresses the challenge of developing a multi-primer approach for DNA metabarcoding analysis to disentangle the most common plants and arthropods resources ingested by field-collected omnivorous predators. The digestion process reduces the likelihood of detecting ingested DNA from gut or whole specimens. One way to improve PCR success in insect diet analyses is to increase the amount of DNA template by pooling individual specimens of the same species. Such pooling has been performed in previous metabarcoding studies to estimate predator diets in bats (*Chalinolobus gouldii* Gray) and birds (*Sialia mexicana* Swainson) (Burgar et al. 2014; Jedlicka et al. 2017), and leads to the detection of most commonly ingested taxa (Mata et al. 2019). This strategy reduces cost and time, like other strategies, as the nested tagging, that have been also used in insect predation studies (Kitson et al. 2019). However, nested tagging can be highly sensitive to cross-contamination between the analysed samples and the control, introducing other biases avoidable with our approach.

Our first objective aimed to determine the effect of a variable sample-pool size (10 or 23 *A. nemoralis*) on the taxonomic coverage. As the number of taxa detected was not significantly different between both sample-pool sizes (Table S3, *Taxonomic coverage* trial), we conducted the *Analysis of field-collected predators* trial by pooling up to 26 individuals together in the same library, in order to save time and cost of the HTS run. Our second objective aimed to compare the performance of using either one or two pairs of primers in the same library.The use of one pair of primers per library is common practice in HTS studies for arthropod (Burgar et al. 2014) and plant-focused studies (Richardson et al. 2015).Here we showed the benefit of using two pairs of arthropod primers together in the same library. On one hand, no significant differences were observed in the number of taxa obtained in both cases in the *Taxonomic coverage* trial. On the other hand, the use of two primer pairs in the same library reduced the number of libraries by half, which consequently decreased the cost, as well as the time needed for the preparation of the libraries. For this reason, both arthropod and both plant pairs of primers were used in one library in the *Analysis of field-collected predator trial*.

When studying the diet of omnivorous species, a multi-primer approach is needed to characterize the full diet, and the choice of the primer pairs is key, because the richness of the taxa obtained depends on it (Hajibabaei et al. 2019). However, some aspects like the taxonomic coverage or the taxonomic resolution of each primer pair used, or their complementarity are not well known, despite their potential impact on the final results (Deagle et al. 2018; Corse et al. 2019).

Considering the three trials of the present study, we observed that the arthropod primer pair mlCOIintF/HC02198 amplified a slightly higher percentage of taxa, around 20% more than ZBJ-ArtF1c/ZBJ-ArtR2c in both batch (Fig. S2 (A)). Both arthropod primer pairs amplify a short fragment within the multicopy COI region, improving the detection of degraded DNA by the digestion process (Agustí et al. 2003). But, even if they amplify fragments in the same region, both primer pairs have different primer binding sites (Fig. S1), which increases the chances to amplify different taxa (Table S3). As suggested by Piñol et al. (2015), this is probably due to the different number of template-mismatches of each arthropod primer pair for each taxon, because a high number of template-mismatches has a negative impact on the amplification efficiency, and reduces the number of amplified taxa. Seven arthropod taxa were detected when using ZBJ-ArtF1c/ZBJ-ArtR2c and 11 with mlCOIintF/HC02198 (Fig. S3 (A)). However, when both pairs of primers were used together, we were able to increase the detection rate up to 14 different arthropod taxa (only four of them were amplified by both primer pairs), showing a higher taxonomic coverage when using both primers instead of only one.

Both plant primer pairs were also selected to have different primer binding sites. The primer pair ITS-S2F/ITS4R amplifies a fragment of the nuclear ITS region, and cA49325/trnL110R of the chloroplast *trnL* region. The first was chosen because it is the most common region to identify mixed pollen loads from insects (Suchan et al. 2019), and the second one because it was recommended for the analysis of degraded DNA (Taberlet et al. 2007). Our results confirmed this statement, as a higher percentage of taxa was amplified with cA49325/trnL110R compared to ITS-S2F/ITS4R in both batch (Fig. S2 (B)), especially in the second batch where DNA was mainly ingested (Table 1). The number of plant taxa detected when considering both trials was 11 for each primer pair (Fig. S3 (B)). However, when using both primer pairs together, we detected 20 different plant taxa, showing a higher taxonomic coverage, as only three taxa were shared by both pairs of primers.

The use of two arthropod primer pairs that generate amplicons of different lengths allow discriminating between those sequences produced by each primer pair. This information was very useful to determine the taxonomic resolution obtained with each primer pairs. Considering all taxa obtained in this study, resolution of both arthropod primer pairs (ZBJ-ArtF1c/ZBJ-ArtR2c and mlCOIintF/HC0219) was mainly to species level (84.31% and 95.96%, respectively) (Fig. S4). On the other hand, resolution of both plant primer pairs (ITS-S2F/ITS4R and cA49325/trnL110R) was mainly to species level (81.82%) and to family level (81.91%), respectively (Fig. S4). These results corroborate those obtained in other studies using the same pairs of primers for metabarcoding and barcoding studies (da Silva et al. 2019; Suchan et al. 2019; Zhu et al. 2019). Such high-level resolution obtained with both arthropod primers and with ITS for plants increases the certainty of the obtained results (Biffi et al. 2017; McInnes et al. 2017; Deagle et al. 2018). Taxonomic resolution should be a factor to consider in the selection of the primer pairs, particularly for plant primers, where the taxonomic resolution is more variable.

### Trophic interactions

In this study, we assumed that plant DNA obtained from whole body extraction of cleaned insects came from their gut contents and corresponded to their diet. On the contrary, plant DNA retrieved from washing solutions is taken to represent visited plants, either from the pollen deposited on their bodies when foraging on them, or from walking on leaves with deposited pollen from anemophilous plants of the surrounding vegetation. We only detected plant DNA from the washed bodies of *R. fulva*, which are larger and hairier than *A. nemoralis*. The fact that we did not detect plant DNA from *A. nemoralis* washing solutions indicates that it may not be necessary to wash such small and glabrous insects. Even if it is well known that anthocorids like *Orius* spp feed on plant resouces in laboratory conditions (Naranjo & Gibson 1996), no plant taxa were detected using the whole specimens either. If their most recent feeding episode was on arthropod prey, that may explain this result. Their small size and their sucking mouthparts, may also explain why no plant food was detected in this species, especially in comparison with *R. fulva*. Plant DNA was identified in only 30% of the analysed individuals of another predatory bug which were present on tomato plants in a greenhouse (Pumariño et al. 2011).

When analyzing the plant taxa ingested by *R. fulva*, we observed that they were all assigned to the Phylum Streptophyta or to a family (Convolvulaceae, Solanaceae, Fabaceae or Poaceae) (Table S3; Fig. 1), being Poaceae and Solanacea the most common detected taxa (Fig. S5 (A)). However, when analyzing their washing solutions, more OTUs were assigned to genera or to species level, possibly because plant DNA from pollen grains attached to the insects’ body is not as degraded as the ingested one. These plant taxa indicate that *R. fulva* forages on a wide range of plants, like *O. europaea, D. glomerata, P. annua, B. vulgaris, Pinus* sp., *Sonchus* sp., one of their family (Asteraceae) and one of their order (Caryophyllales) (Table S3; Fig. 1). This diet is much more diverse than the single plant species cited by Rodwell et al. (2018), *Heracleum sphondylium* L (Apiaceae). The detected plant taxa can be present in ground covers of peach crops, field margins or alfalfa crops in the area of study (Juarez-Escario et al. 2010; Juarez-Escario et al.,2018; Solé-Senan et al. 2018), and some of them, like *D. glomerata, P. annua* and *M. sativa* belong to families that were also detected by ingestion (Poaceae and Fabaceae), which may indicate that their body was in contact with pollen from tassels or flowers while consuming it.

In the *Analysis of field-collected predators* trial, we have also demonstrated the efficacy of this multi-primer approach to detect and identify arthropods ingested by both predator species (Table S3; Fig. 1). Even if previous literature cites *R. fulva* as predator of some insects, such as aphids (Pons & Eizaguirre, 2000; Rodwell et al. 2018), our results indicate that this predator also consumed *N. graminicola*, because it was detected in 25% of the analysed *R. fulva* sample-pools (Fig. S5 (B)). *Nisius graminicola* is cited as an important pest of several summer crops in Italy, including peaches (Blando & Mineo, 2005). In Spain, another species of the same genus, *N. ericae*, has been described as secondary pest in peaches (Del Rivero & García-Marí, 1983), thus suggesting the potential of *R. fulva* as biocontrol agent. Our results also show that intraguild predation (IGP) by *R. fulva* on Coccinellidae and *C. livida*, is a very common trophic interaction (Fig. S5 (B)). It is well known that IGP is widespread in agroecosystems (Lucas and Rosenheim, 2011), and HTS has been successful at demonstrating IGP for example in field-collected predators in lettuce (Gomez-Polo et al. 2015, 2016). The IGP observed here should be further studied in order to know whether it could have a negative effect on the biological control of peach pests, because some coccinellids such as *C. septempunctata* or *Stethorus punctillum* (Weise) are efficient biocontrol agents in peach orchards (Trandafirescu et al. 2004; Biddinger et al. 2009).

*Anthocoris nemoralis* is a well-known biocontrol agent in fruit orchards, particularly of the pear psylla (Solomon et al. 2000; Agustí et al. 2003). Our results indicate that this species is in fact a polyphagous predator, since its most common prey in our study were two very important peach pests, the green peach aphid *M. persicae*, and the peach moth *G. molesta* (Fig. S5; Table S3; Fig. 1), information unknown until now. This predator also fed on *D. lycii*, an hemipteran species which is oligophagous on *Lycium* plants (Solanaceae). Since *Lycium europaeum* L. is planted in hedges to separate agricultural plots in the study area, it can be assumed that *A. nemoralis* must have moved from peach to those hedges to feed on this particular prey species and then back to the crop were it was collected. This result demonstrates how HTS analysis could also be used as a tool to understand predator movement, in this case from the peach crop to the surrounding vegetation and backwards. Finally, we also detected IGP in *A. nemoralis* (Fig. 1), which fed on several species coccinellid in the genus *Orius*. These included *O. conglobata*, a very common species in urban landscapes (Lumbierres et al., 2018), and *O. laevigatus*, a known biocontrol agent in vegetables (Gomez-Polo et al. 2015, 2016). The latter trophic interaction should be also taken in consideration in further biological control studies.

Four arthropod taxa were also detected in the diet of *A. nemoralis* analysed in the *Taxonomic coverage* trial (Table S3; Fig. 1), reinforcing its role as generalist predator. One of them was *T. fuscipennis*, which damages peaches during ripening (Tavella et al. 2006). Also detected, the subfamily *Cecidomyiinae* includes some predator species and some gall-producing pests in forestry and horticulture (Kolesik, 2014). The other two prey taxa were in the genus *Orius* and in the family Coccinellidae, which are predators known to be present in both crops in the area of study (Trandafirescu et al. 2004; Pons et al. 2009; Aparicio et al. 2020). Our results reinforce the role of *A. nemoralis* as potential biological control agent, which should be considered in further studies in peach orchards and alfalfa crops. This is especially important in the study area where both crops coexist and the movement of insects between them is very likely.

In this study, we have detected arthropod and plant resources ingested by two insect predators present in a peach crop by HTS analysis using a multi-primer approach. We have demonstrated that pooling predators in groups of 10 or 23 individuals has no significant influence on the analysis of their diet when analysed this way. We also showed that the use of two primer pairs improves the detection of ingested taxa, with an increased number of arthropod and plant taxa. Finally, we have shown that washing predators prior to HTS analysis is particularly needed for large insects with hairy structures, but may not be useful for small and glabrous ones. The developed multi-primer approach reduces time and cost of the HTS analysis and shows both expected and unexpected trophic relationships. The description of the most common trophic interactions by HTS with multi-primer approach could lead to an improvement of the biological control of pest species in agroecosystems, contributing to a more sustainable agriculture. The detection of a wider than expected range of ingested arthropod and plant items highlights the importance of keeping a diverse landscape composition in order to enhance the conservation of biological control agents in crops.

## Acknowledgements

The authors would like to thank Lorena Gallardo and Angels Tudó for their technical assistance in field collecting samples and laboratory procedures. The landowners of the crop plot are also acknowledged for allowing us to access to their fields. This research was funded by the Spanish Ministry of Economy, Industry and Competitiveness (grant AGL2014-53970-C2-2-R) and by the CERCA Programme (Centres de Recerca de Catalunya) of the Generalitat de Catalunya. Ivan Batuecas was funded by the grant BES-2015-075700 from the Ministry of Science, Innovation and Universities.

