## supplemental_materials for "Development of a multi-primer metabarcoding approach to understand trophic interactions in agroecosystems"

**Table S1.** Arthropod (ZBJ-ArtF1c/ZBJ-ArtR2c and mlCOIintF/HC02198) and plant (ITS-S2F/ITS4R and cA49325/trnL110R) primer pairs used in this study, indicating the sequence of each forward (F) and reverse (R) primer and the length of the amplified fragment.

**Resources detected**

**Predator analysed**

|  | Sequence 5´- 3´(F) | Sequence 5´- 3´(R) | Reference | Fragment (bp) |
| --- | --- | --- | --- | --- |
| ZBJ-ArtF1c/ZBJ-ArtR2c | AGATATTGGAACWTTATATTTTATTTTTGG | WACTAATCAATTWCCAAATCCTCC | Zeale et al. 2011 | 157 |
| mlCOIintF/HC02198 | GGWACWGGWTGAACWGTWTAYCCYCC | TAAACTTCAGGGTGACCAAAAAATCA | Leray et al. 2013/Folmer et al. 1994 | 313 |
| ITS-S2F/ITS4R | ATGCGATACTTGGTGTGAAT | TCCTCCGCTTATTGATATGC | Chen et al. 2010/White et al. 1990 | 350 |
| cA49325/trnL110R | CGAAATCGGTAGACGCTACG | GATTTGGCTCAGGATTGCCC | Taberlet et al. 2007/Borsch et al. 2003 | 80 |

**Table S2.** Relative read abundance (RRA) obtained from each arthropod and plant primer pairs in each library (L) in the three trials included in the study: (1) *Taxonomic coverage*; (2) *Plant primer resolution*; and (3) *Analysis of field-collected predators*. 3A corresponds to arthropods, 3B to plants, and 3C to washing solutions. Those percentages eliminated from the analysis for not reaching the 1% threshold are shown in bold. Art1= ZBJ-ArtF1c/ZBJ-ArtR2c; Art2= mlCOIintF/HC02198; Pl1= ITS-S2F/ITS4R; Pl2= cA49325/trnL110R; NA= Not amplified.

**(1)**

|  | **L1** | **L2** | **L3** | | **L4** | **L5** | **L6** | |
| --- | --- | --- | --- | --- | --- | --- | --- | --- |
|  | **Art1** | **Art2** | **Art1** | **Art2** | **Art1** | **Art2** | **Art1** | **Art2** |
| *Orius* | - | - | - | - | - | 20,44 | **-** | 18,15 |
| *Orius laevigatus* | - | - | - | - | 22,72 | **0,03** | 22,19 | **-** |
| Aphididae | - | - | 18,17 | **-** | 28,69 | **-** | 28,74 | **-** |
| *Myzus persicae* | - | 21,92 | - | 27,16 | - | 31,71 | **-** | 35,67 |
| *Diaphorina lycii* | - | 1,57 | - | 1,33 | - | 1,36 | - | 1,09 |
| Coccinellidae | **0,02** | - | - | - | **0,35** | **-** | **0,35** | **-** |
| Cecidomyiinae | 2,18 | **-** | 1,93 | **-** | 3,91 | **-** | 3,67 | **-** |
| *Grapholita molesta* | 97,80 | 76,51 | 79,90 | 71,52 | 44,06 | 45,15 | 44,77 | 43,93 |
| *Thrips fuscipennis* | - | - | - | - | **-** | 1,32 | **-** | 1,16 |
| *Sitona discoideus* | - | - | - | - | **0,27** | **-** | **0,28** | **-** |

**(2)**

|  | **L7** | | **L8** | | **L9** | | **L10** | |
| --- | --- | --- | --- | --- | --- | --- | --- | --- |
|  | **Pl1** | **Pl2** | **Pl1** | **Pl2** | **Pl1** | **Pl2** | **Pl1** | **Pl2** |
| Streptophyta | **-** | **0,17** | **-** | **-** | **-** | **-** | - | - |
| Asteraceae | **-** | 39,94 | **-** | **-** | **-** | **-** | - | - |
| *Picris echioides* | 16,13 | **-** | **-** | **-** | **-** | **-** | - | - |
| Convolvulaceae | **-** | **0,64** | **-** | **-** | **-** | **-** | - | - |
| *Convolvulus arvensis* | 15,36 | **-** | **-** | **-** | **-** | **-** | - | - |
| Fabaceae | **-** | 2,29 | **-** | **-** | **-** | 99,97 | - | 99,94 |
| *Medicago_sativa* | 23,75 | **-** | **-** | **-** | 100 | **-** | 100 | **0,05** |
| *Trifolium* | **-** | 2,79 | **-** | **-** | **-** | **-** | - | - |
| *Setaria* | 5,11 | 34,19 | **-** | **-** | **-** | **-** | - | - |
| Rosaceae | **-** | 20 | **-** | 100 | **-** | **0,03** | - | **0,004** |
| *Prunus persica* | 39,65 | - | 100 | - | **-** | - | - |  |

**(3A)**

|  | **L11** | | **L13** | | **L15** | | **L17** | | **L19** | | **L21** | |
| --- | --- | --- | --- | --- | --- | --- | --- | --- | --- | --- | --- | --- |
|  | **Art1** | **Art2** | **Art1** | **Art2** | **Art1** | **Art2** | **Art1** | **Art2** | **Art1** | **Art2** | **Art1** | **Art2** |
| *Orius* | **-** | **-** | **-** | **-** | **-** | **-** | NA | **-** | **-** | **-** | **-** | 18,32 |
| *Orius laevigatus* | **-** | **-** | **-** | **-** | **-** | **-** | NA | **-** | **-** | **-** | 21,09 | **-** |
| Aphididae | **-** | **-** | **-** | **-** | **-** | **-** | NA | **-** | 100 | - | 22,54 | **-** |
| *Myzus persicae* | **-** | **-** | **-** | **-** | **-** | **-** | NA | **-** | **-** | 24,39 | **-** | 24,29 |
| *Nysius graminicola* | **0,06** | - | 50 | 42,11 | **-** | **-** | NA | **-** | **-** | - | **-** | - |
| *Diaphorina lycii* | **-** | - | - | - | **-** | **-** | NA | **-** | **-** | 37,80 | - | 1,90 |
| Coccinelidae | **-** | - | - | 57,89 | **-** | **0,31** | NA | 100 | **-** | - | **0,70** | - |
| *Oenopia conglobata* | **-** | - | - | **-** | - | - | NA | **-** | **-** | 37,80 | - | 5,70 |
| *Cantharis_livida* | 99,94 | 100 | 50 | **-** | 100 | 99,69 | NA | **-** | **-** | **-** | **-** | **-** |
| Cecidomyiinae | **-** | - | - | **-** | - | - | NA | **-** | **-** | **-** | 5,00 | - |
| *Grapholita molesta* | **-** | - | - | **-** | - | - | NA | **-** | - | - | 50,68 | 39,76 |
| *Thrips fuscipennis* | **-** | - | - | **-** | - | - | NA | **-** | - | - |  | 10,04 |

**(3B)**

|  | **L12** | | **L14** | | **L16** | | **L18** | | **L20** | | **L22** | |
| --- | --- | --- | --- | --- | --- | --- | --- | --- | --- | --- | --- | --- |
|  | **Pl1** | **Pl2** | **Pl1** | **Pl2** | **Pl1** | **Pl2** | **Pl1** | **Pl2** | **Pl1** | **Pl2** | **Pl1** | **Pl2** |
| Streptophyta | NA | - | - | 13,04 | NA | - | NA | 22,29 | NA | NA | NA | NA |
| Convolvulaceae | NA | - | - | 56,52 | NA | - | NA | 33,76 | NA | NA | NA | NA |
| Solanaceae | NA | 37,65 | - | 6,52 | NA | - | NA | 35,03 | NA | NA | NA | NA |
| Fabaceae | NA | 62,35 | - | **-** | NA | 47,37 | NA | - | NA | NA | NA | NA |
| Poaceae | NA | - | 100 | 23,91 | NA | 52,63 | NA | 8,92 | NA | NA | NA | NA |

**(3C)**

| **Libraries** | **L23** | | **L24** | | **L25** | | **L26** | | **L27** | |
| --- | --- | --- | --- | --- | --- | --- | --- | --- | --- | --- |
| **Primers pairs** | **Pl1** | **Pl2** | **Pl1** | **Pl2** | **Pl1** | **Pl2** | **Pl1** | **Pl2** | **Pl1** | **Pl2** |
| Streptophyta | - | 63,95 | - | 6,11 | - | 86,48 | NA | NA | NA | NA |
| Asteraceae | - | 7,08 | - | 81,22 | - | 4,44 | NA | NA | NA | NA |
| *Sonchus* | 12,75 | - | - | - | 28,41 | - | NA | NA | NA | NA |
| *Medicago sativa* | 1,24 | - | - | - | 2,21 | - | NA | NA | NA | NA |
| *Olea europaea* | 5,17 | - | - | - | 6,09 | 1,25 | NA | NA | NA | NA |
| *Pinus* | - | 23,01 | - | - | - | - | NA | NA | NA | NA |
| Poaceae | 28,33 | 4,43 | - | 9,61 | 41,70 | 7 | NA | NA | NA | NA |
| *Dactylis glomerata* | 4,30 | - | - | - | 4,06 | - | NA | NA | NA | NA |
| *Poa annua* | 19,88 | - | - | - | 3,69 | - | NA | NA | NA | NA |
| Caryophyllales | - | 1,53 | - | 3,06 | - | **0,83** | NA | NA | NA | NA |
| *Beta vulgaris* | 28,33 | - | - | - | 13,84 | - | NA | NA | NA | NA |

**Table S3.** Taxa obtained and primer pairs used in each trial and from each library. NT= not tested; NAmp = not amplified

| **Trial** | **Species/sample** | **# of individuals/**  **Sample size** | **Library** | **Assigned taxa with each arthropod primer pair** | | **Assigned taxa with each plant primer pair** | |
| --- | --- | --- | --- | --- | --- | --- | --- |
|  |  |  |  | **ZBJ-ArtF1c/ZBJ-ArtR2c** | **mlCOIintF/HC02198** | **ITS-S2F/ITS4R** | **CA49325/trnL110R** |
| *Taxonomic coverage* | *Anthocoris nemoralis* | 10 | L1 | Aphididae  Cecidomyiinae  *Grapholita molesta* | NT | NT | NT |
|  |  |  | L2 | NT | *Anthocoris nemoralis*  *Myzus persicae*  *Diaphorina lycii*  *Grapholita molesta* | NT | NT |
|  |  |  | L3 | Aphididae  Cecidomyiinae  *Grapholita molesta* | *Anthocoris nemoralis*  *Myzus persicae*  *Diaphorina lycii*  *Grapholita molesta* | NT | NT |
|  |  | 23 | L4 | *Orius laevigatus*  Aphididae  Cecidomyiinae  *Grapholita molesta* | NT | NT | NT |
|  |  |  | L5 | NT | *Anthocoris nemoralis*  *Orius*  *Myzus persicae*  *Diaphorina lycii*  *Grapholita molesta*  *Thrips fuscipennis* | NT | NT |
|  |  |  | L6 | *Orius laevigatus*  Aphididae  Cecidomyiinae  *Grapholita molesta* | *Anthocoris nemoralis*  *Orius*  *Myzus persicae*  *Diaphorina lycii*  *Grapholita molesta*  *Thrips fuscipennis* | NT | NT |
| *Plant primer*  *resolution* | *Medicago sativa*  *Prunus persica*  *Convulvulus arvensis*  *Picris echiodies*  *Setaria pumila* | 1 cm^2^ | L7 | NT | NT | *Picris echioides*  *Convolvulus arvensis*  *Medicago sativa*  *Setaria*  *Prunus persica* | Streptophyta  Asteraceae  Convolvulaceae  Fabaceae  Trifolium  *Setaria*  Rosaceae |
|  | *Prunus persica* | 1 cm^2^ | L8 | NT | NT | *Prunus persica* | *Rosaceae* |
|  | *Medicago sativa* | 1 cm^2^ | L9 | NT | NT | *Medicago sativa* | Fabaceae  *Rosaceae* |
|  | *Medicago sativa* | 1 cm^2^ | L10 | NT | NT | *Medicago sativa* | Fabaceae  Rosaceae |
| *Analysis of*  *field-collected*  *predators* | *Rhagonycha fulva* | 26 | L11 | *Rhagonycha fulva*  *Cantharis lívida* | *Rhagonycha fulva*  *Cantharis lívida* | NT | NT |
|  |  |  | L12 | NT | NT | NAmp | Fabaceae  Solanaceae |
|  |  | 24 | L13 | *Rhagonycha fulva Nysius graminicola*  *Cantharis lívida* | *Rhagonycha fulva*  *Nysius graminicola* Coccinellidae | NT | NT |
|  |  |  | L14 | NT | NT | Poaceae | Streptophyta  Convolvulaceae  Solanaceae  Poaceae |
|  |  | 23 | L15 | *Rhagonycha fulva Cantharis lívida* | *Rhagonycha fulva Cantharis lívida* | NT | NT |
|  |  |  | L16 | NT | NT | NAmp | Fabaceae  Poaceae |
|  |  | 5 | L17 | *Rhagonycha fulva* | *Rhagonycha fulva* Coccinellidae | NT | NT |
|  |  |  | L18 | NT | NT | NAmp | Streptophyta  Convolvulaceae  Fabaceae  Poaceae |
|  | *Anthocoris nemoralis* | 26 | L19 | Aphididae | *Diaphorina lycii*  *Myzus persicae*  *Oenopia conglobata* *Anthocoris nemoralis* | NT | NT |
|  |  |  | L20 | NT | NT | NAmp | NAmp |
|  |  | 16 | L21 | Aphididae  *Grapholita molesta*  *Orius laevigatus*  Cecidomyiinae | *Anthocoris nemoralis*  *Diaphorina lycii*  *Grapholita molesta*  *Myzus persicae*  *Oenopia conglobata*  *Orius*  *Thrips fuscipennis* | NT | NT |
|  |  |  | L22 | NT | NT | NAmp | NAmp |
|  | *Rhagonycha fulva*  *washing solution* | 26 | L23 | NT | NT | *Sonchus*  *Medicago sativa*  *Olea europaea*  *Dactylis glomerata*  Poaceae  Poa annua  Beta vulgaris | Streptophyta  *Asteraceae*  *Pinus*  Poaceae  Caryophyllales |
|  |  | 24 | L24 | NT | NT | NAmp | NAmp |
|  |  | 23 | L25 | NT | NT | *Sonchus*  *Medicago_sativa*  *Olea_europaea*  *Dactylis_glomerata*  Poaceae  *Poa annua*  *Beta vulgaris* | Streptophyta  Asteraceae  Poaceae  Caryophyllale*s* |
|  |  | 5 | L26 | NT | NT | NAmp | NAmp |
|  | *Anthocoris nemoralis*  *washing solution* | 42 | L27 | NT | NT | NAmp | NAmp |

**Table S4.** Comparison of the number of arthropod taxa obtained in each library in the *Taxonomic coverage* trial with*:* (A) either 10 or 23 *Anthocoris nemoralis* specimens; (B) either one or two pairs of primers.

**(A)**

| **Library** | **# of individuals per library** |  | **# of arthropod taxa** |
| --- | --- | --- | --- |
| L1 | 10 |  | 2 |
| L2 | 10 |  | 4 |
| L3 | 10 |  | 7 |
| L4 | 23 |  | 4 |
| L5 | 23 |  | 6 |
| L6 | 23 |  | 10 |

**(B)**

|  | **Library** | **# of primers per library** |  | **# of arthropod taxa** |
| --- | --- | --- | --- | --- |
|  | L3 | 2 |  | 7 |
|  | L6 | 2 |  | 10 |
|  | L1, L2 | 1 |  | 6 |
|  | L4, L5 | 1 |  | 10 |

**Table S5.** Number of template-mismatches of each arthropod pair of primers with each taxon amplified. NS = no sequence present in the databases at the moment of the analysis.

|  | **mlCOIintT/HC02198** | | **ZBJ-ArtF1c/ ZBJ-ArtR2c** | |
| --- | --- | --- | --- | --- |
| **Taxon** | **Forward primer (mlCOIintT)** | **Reverse primer (HC02198)** | **Forward primer (ZBJ-ArtF1c)** | **Reverse primer (ZBJ-ArtR2c)** |
| *Anthocoris nemoralis* | 0 | 0 | 4 | 5 |
| Aphididae | 1 | NS | 3 | 2 |
| *Cantharis livida* | 0 | NS | 1 | 2 |
| Cecidomyiinae | 0 | NS | 1 | 1 |
| Coccinellidae | 1 | 3 | 3 | 3 |
| *Diaphorina lycii* | 1 | 3 | 10 | 9 |
| *Grapholita molesta* | 1 | 3 | 0 | 0 |
| *Myzus persicae* | 2 | 2 | 3 | 3 |
| *Nysius graminicola* | 1 | 2 | 0 | 0 |
| *Oenopia conglobata* | 0 | NS | 12 | 2 |
| *Orius* | 2 | 4 | 3 | 3 |
| *Orius laevigatus* | 0 | NS | 2 | 0 |
| *Rhagonycha fulva* | 1 | NS | 2 | 2 |
| *Thrips fuscipennis* | 1 | NS | 8 | 12 |

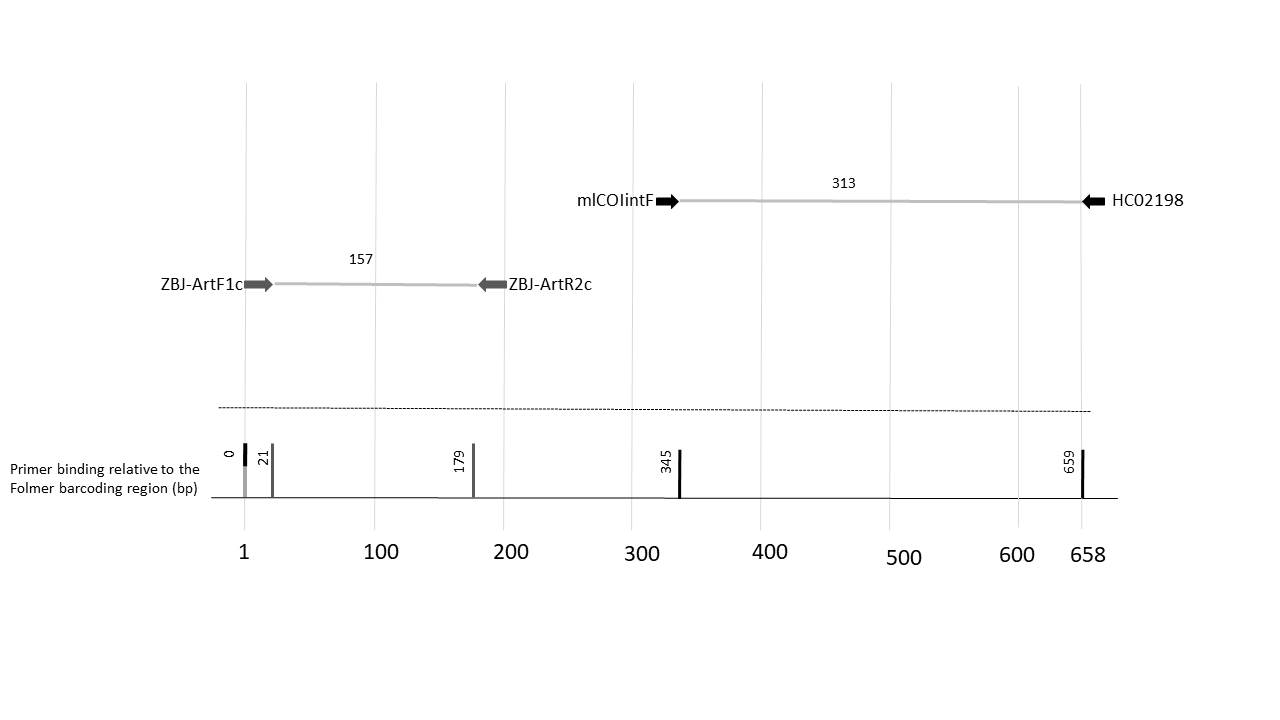
**Figure S1.** Location of the COI primer pairs tested in the present study.

**Figure S2.** Percentage of taxa obtained for each pair of arthropod (A) and plant (B) primers amplified in all libraries, in each HTS batch.

40%

60%

(A)

Batch 1

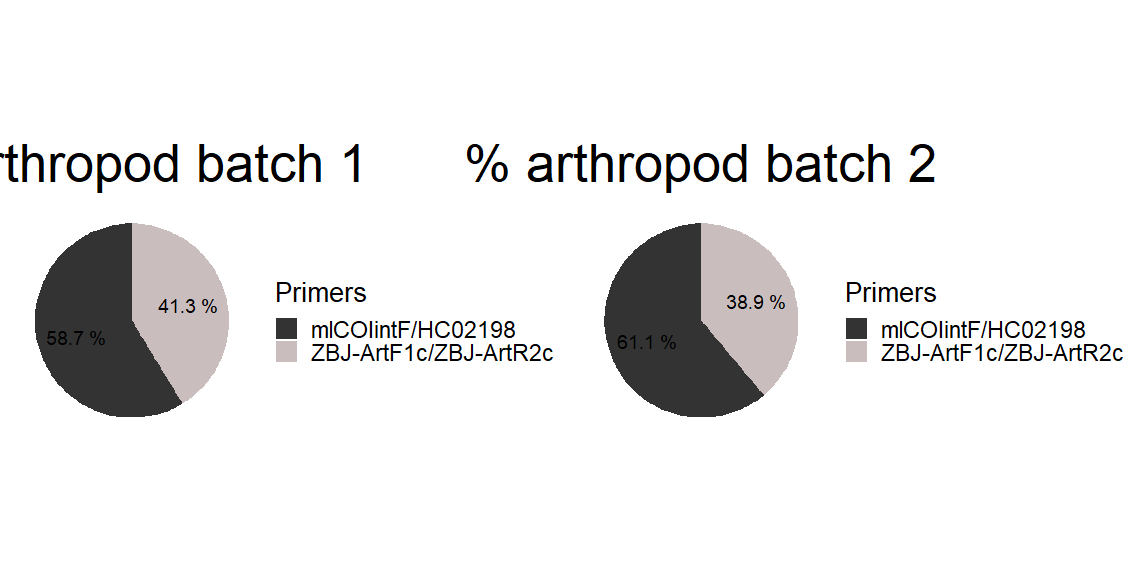

ZBJ-ArtF1c/ZBJ-ArtR2c

mlCOIintF/HC02198

Batch 2

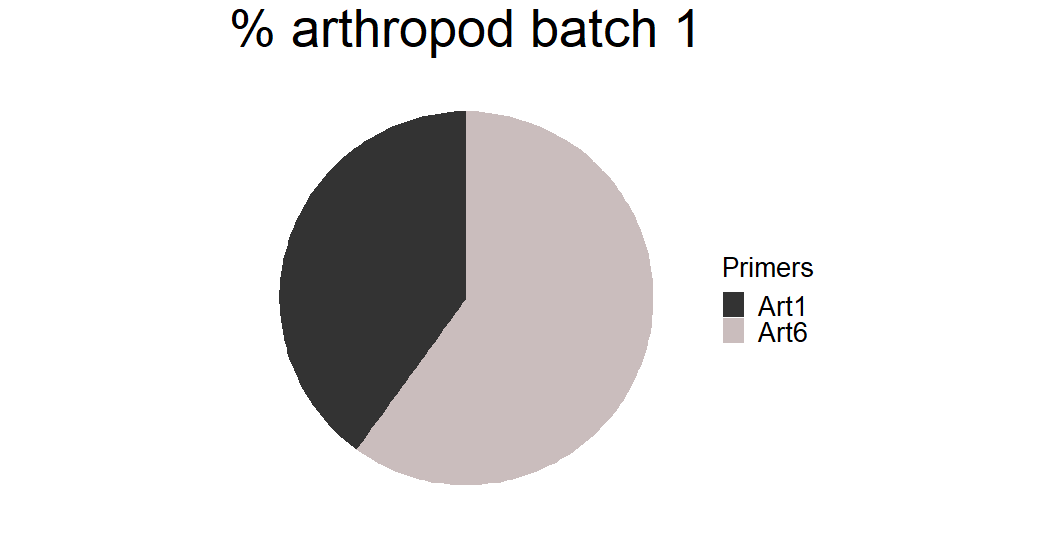

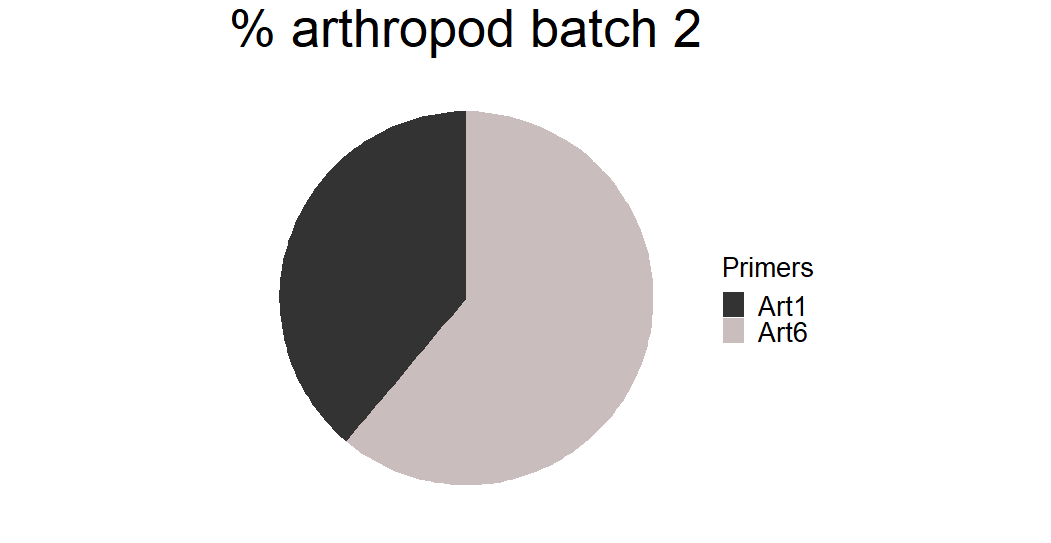

61.12%

38.88%

(B)

58.34%

41.66%

Batch 1

Batch 2

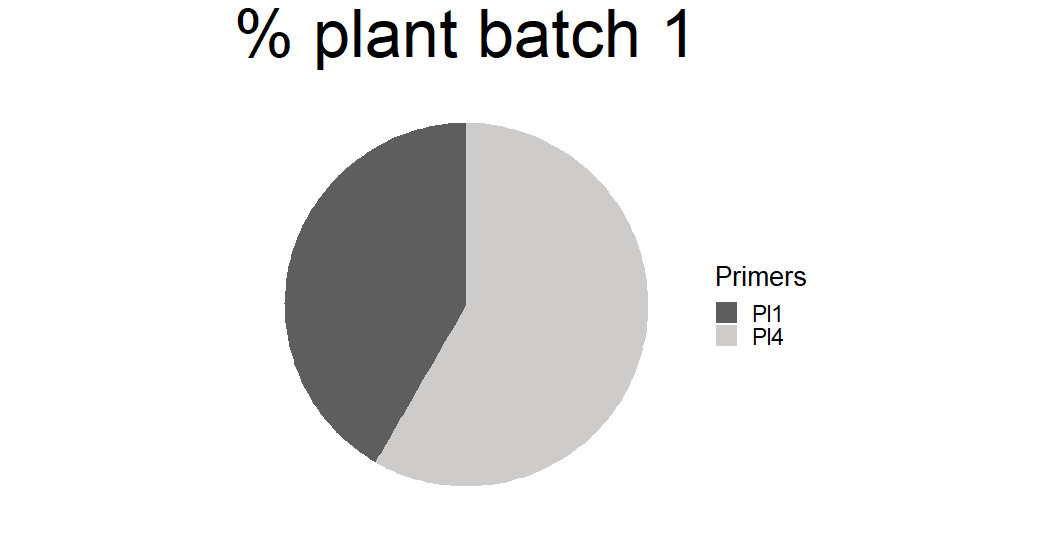

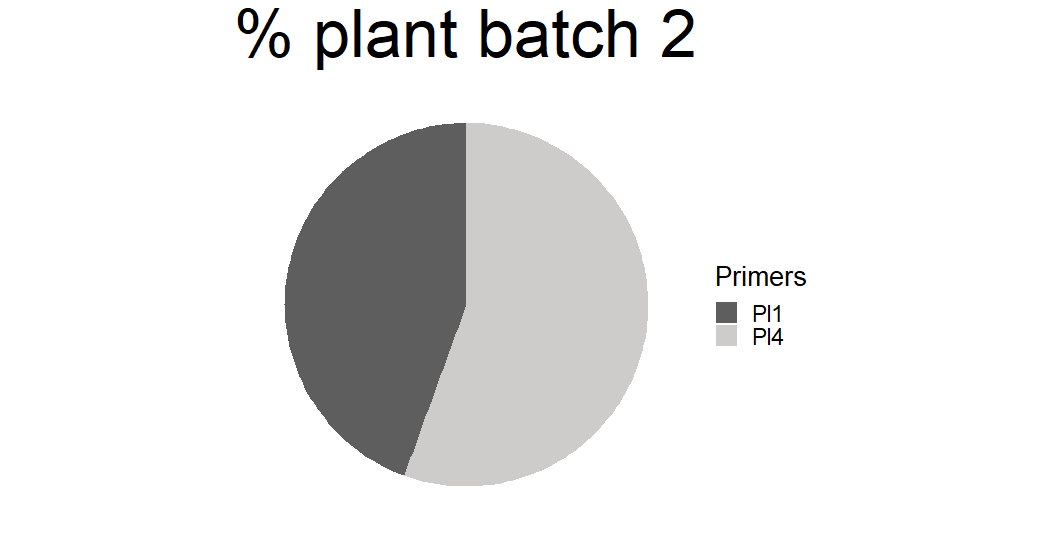

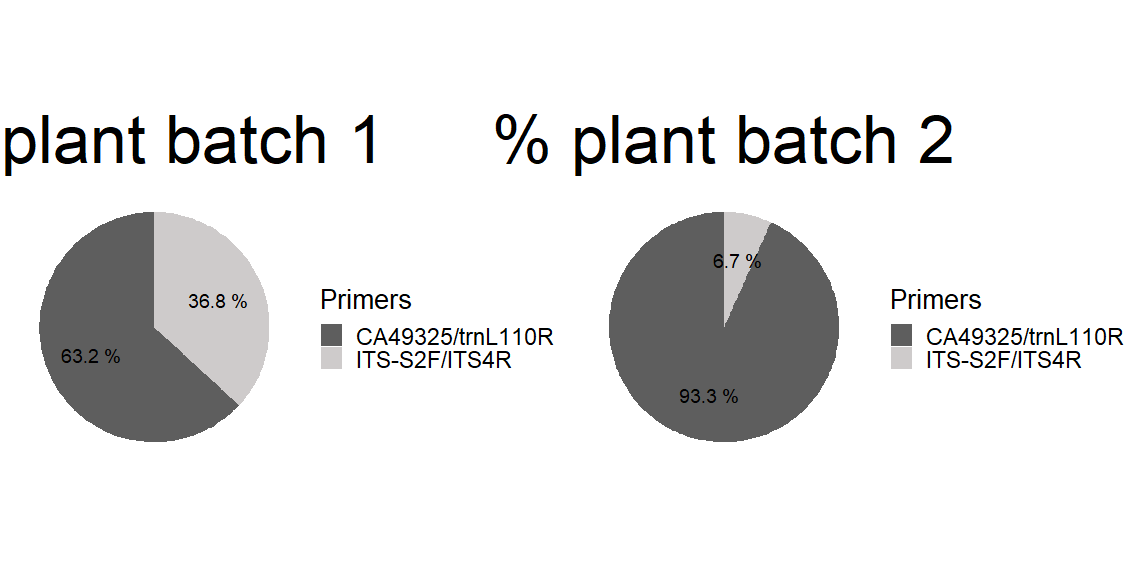

CA49325/trnL110R

ITS-S2F/ITS4R

55.56%

44.44%

**Figure S3.** Representation by Venn´s diagrams of the artropod and plant taxa obtained by HTS (including all trials) with each primer pair. Numbers in each circle indicate the number of taxa amplified by each primer and how many are shared by both pairs of primers (overlaping area). (A) artropod primers; (B) plant primers. n = Number of taxa obtained with each primer pair.

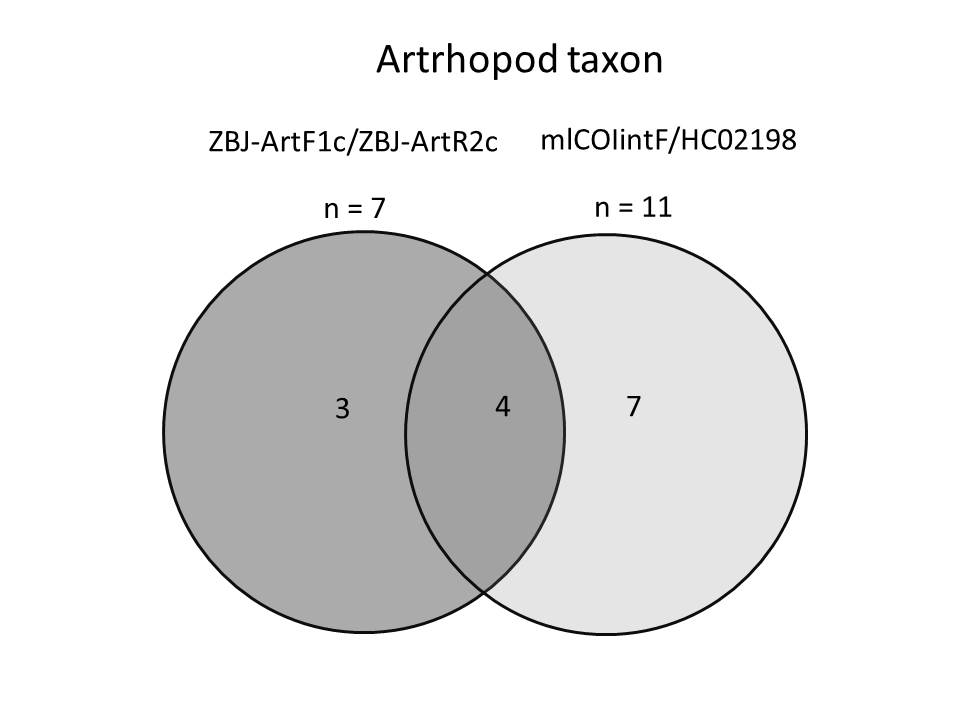
(A)

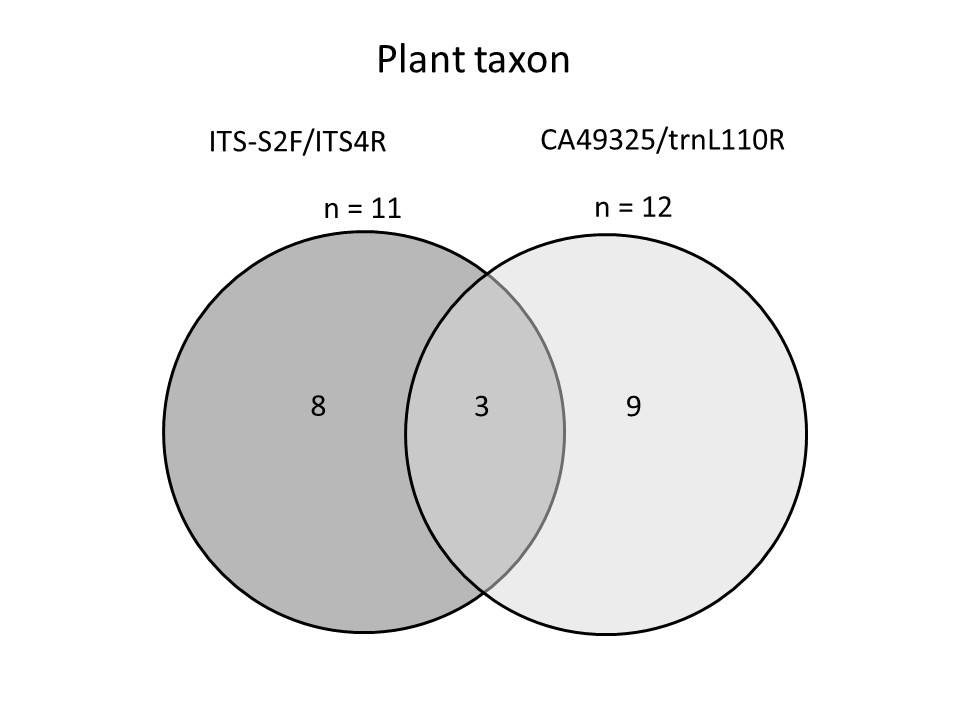
(B)

**Figure S4.** Accuracy of the taxonomic assignation according to the primer pair used. Data is presented as the percentage OTUs assigned to each taxonomic level.

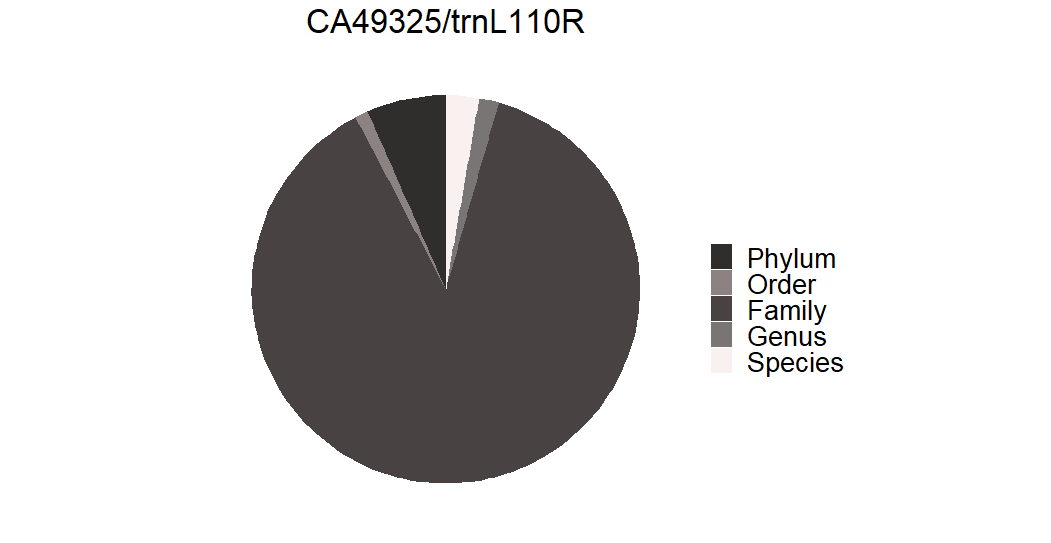

CA49325/trnL110R

1.09%

6.59%

2.74%

87.91%

1.64%

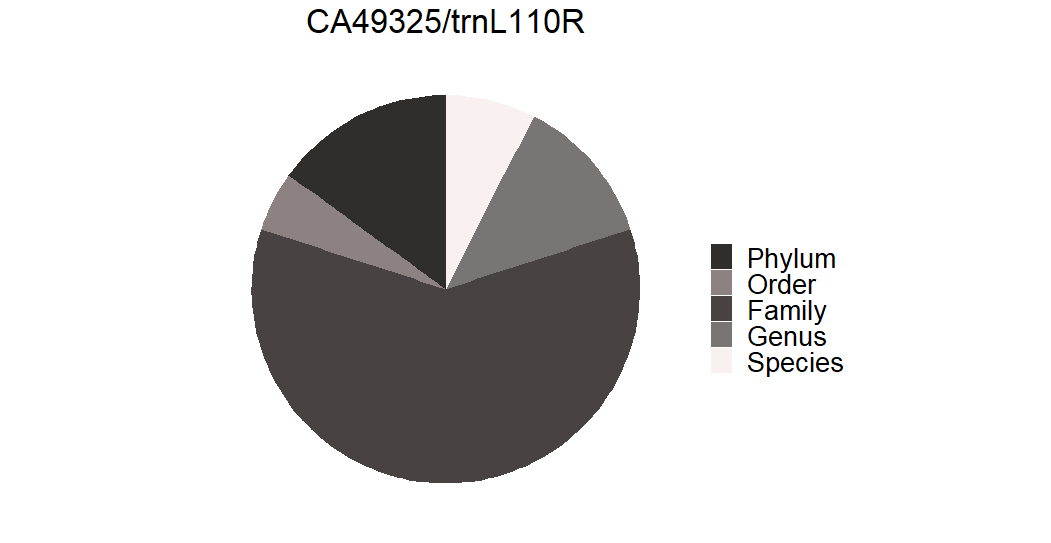

mlCOIintF/HC02198

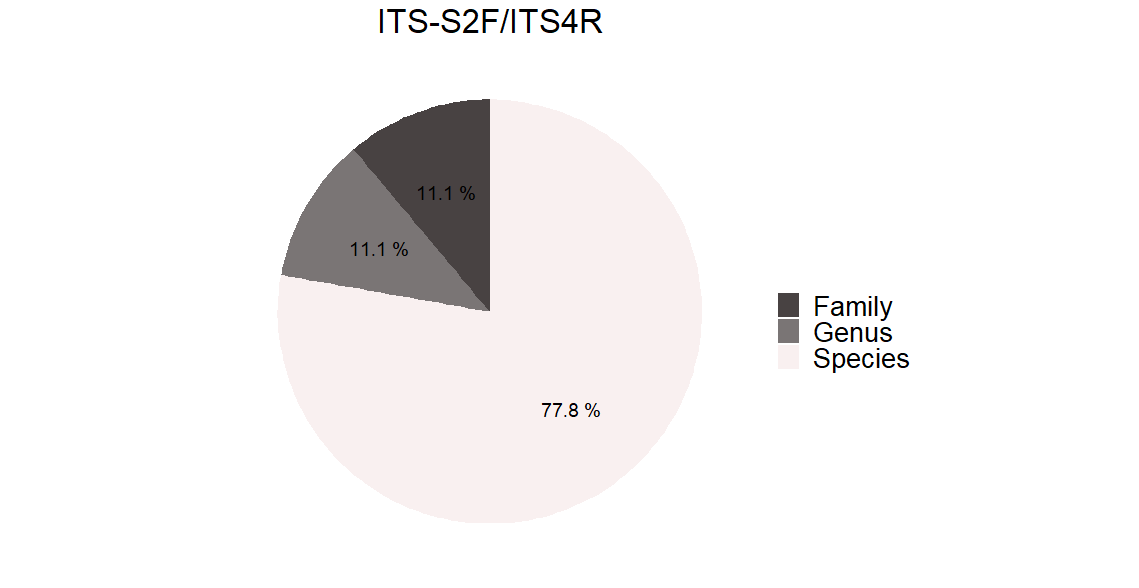

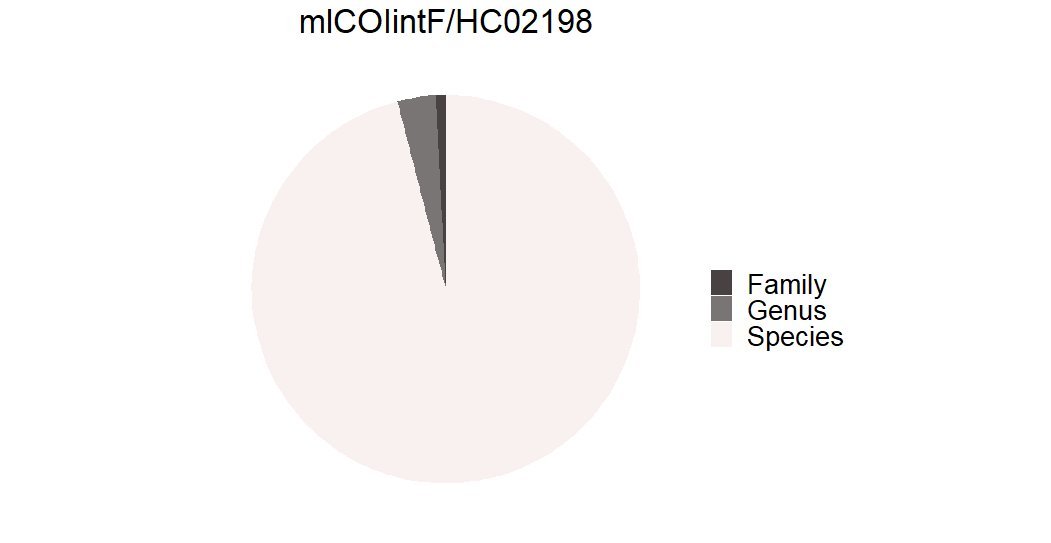

95.96%

0.82%

3.22%

ZBJ-ArtF1c/ZBJ-ArtR2c

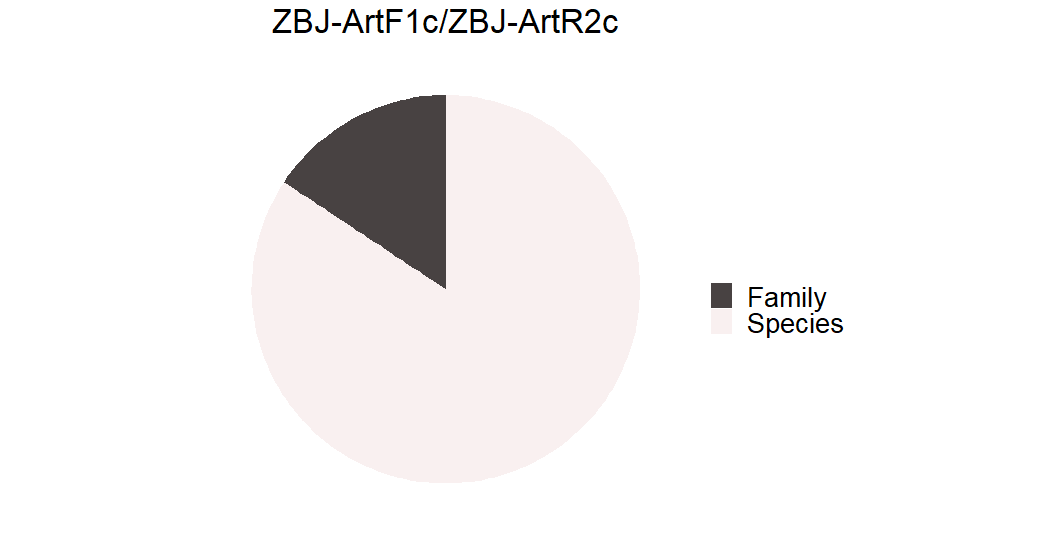

15.69%

84.31%

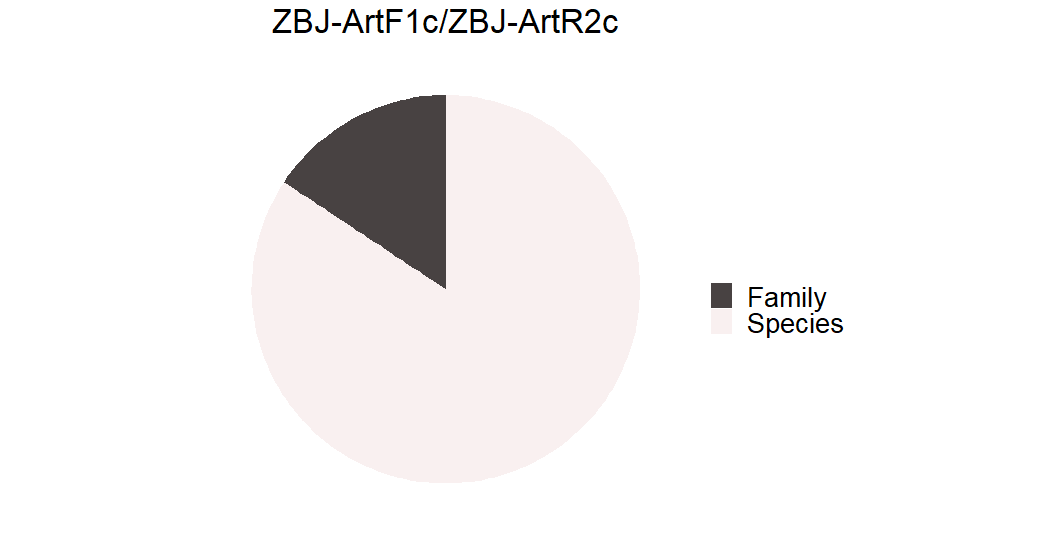

ITS-S2F/ITS4R

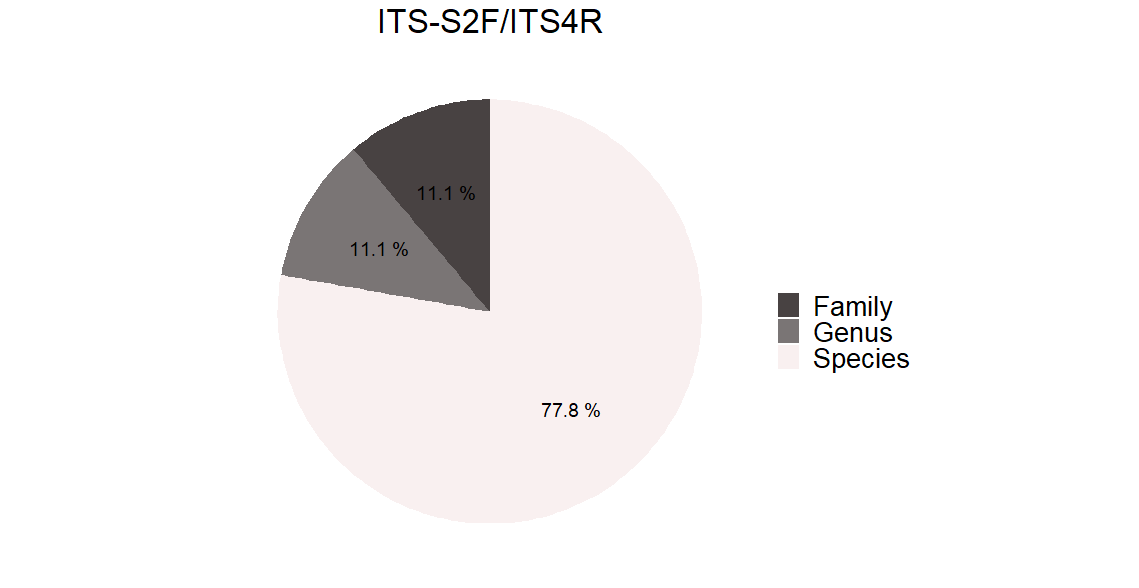

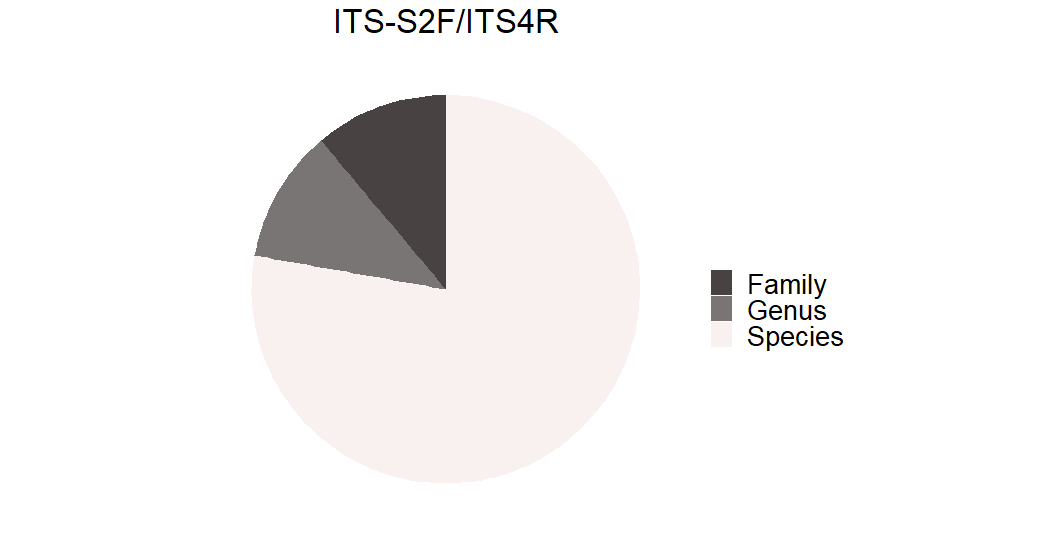

9.09%

9.09%

81.82%

**Figure S5.** Percentage of frequency of occurrence (FOO%) of the obtained taxa items: (A) plant consumed by *R. fulva* and detected in *R. fulva* washing solutions; and (B) arthropod consumed by *A. nemoralis* and *R. fulva*.

(A)

Washing solutions of *R.fulva*

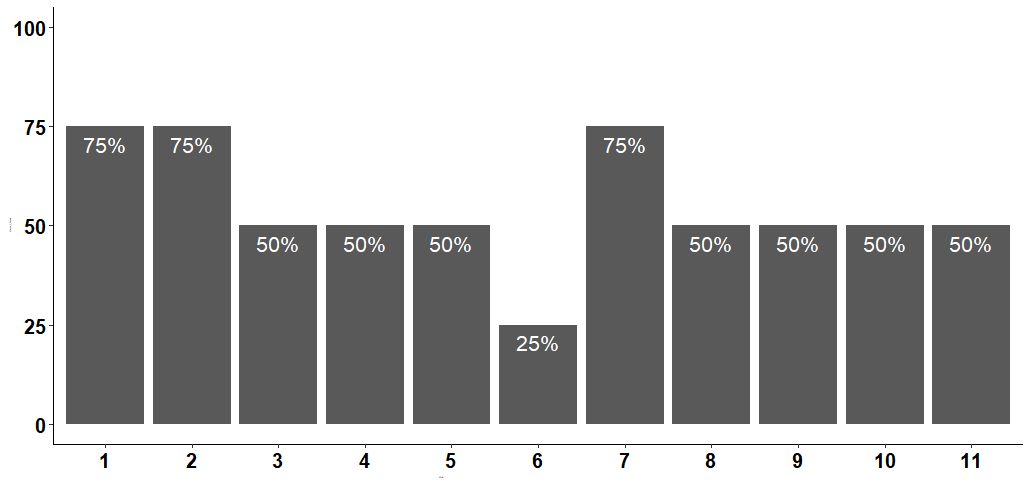

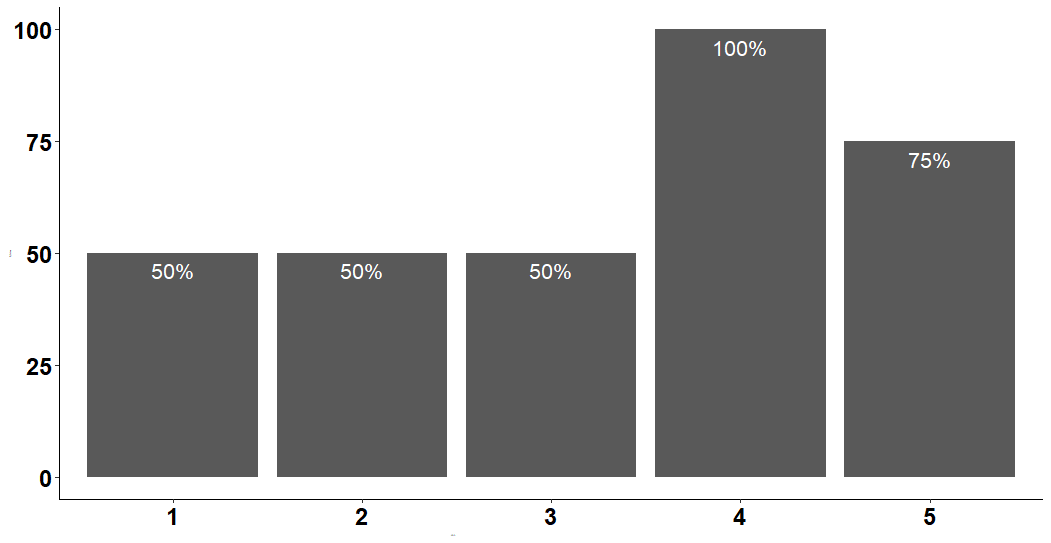
 *R. fulva*

1-Strepthopyta

2-Convolculaceae

3-*Fabaceae*

4-Poaceae

5-Solanaceae

**Taxa**

**%**

1-Strepthopyta

2-Asteraceae

3-*Sonchus*

*4-Medicago sativa*

*5-Olea europaea*

6-*Pinus*

7-*Poaceae*

8-*Dactylys glomerata*

9-*Poa annua*

10-Caryophyllales

11-*Beta vulgaris*

**Taxa**

**%**

**Taxa**

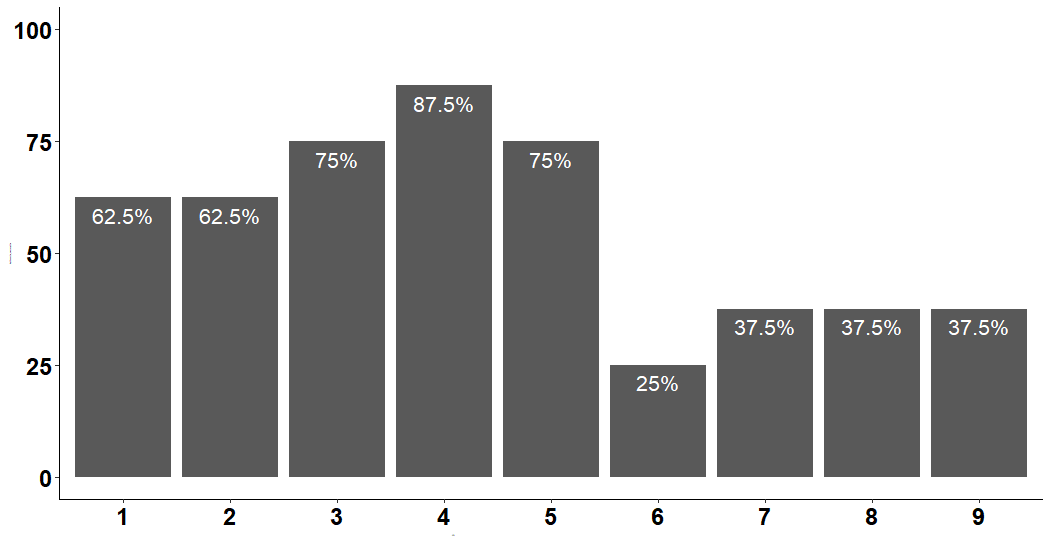
**(B)**

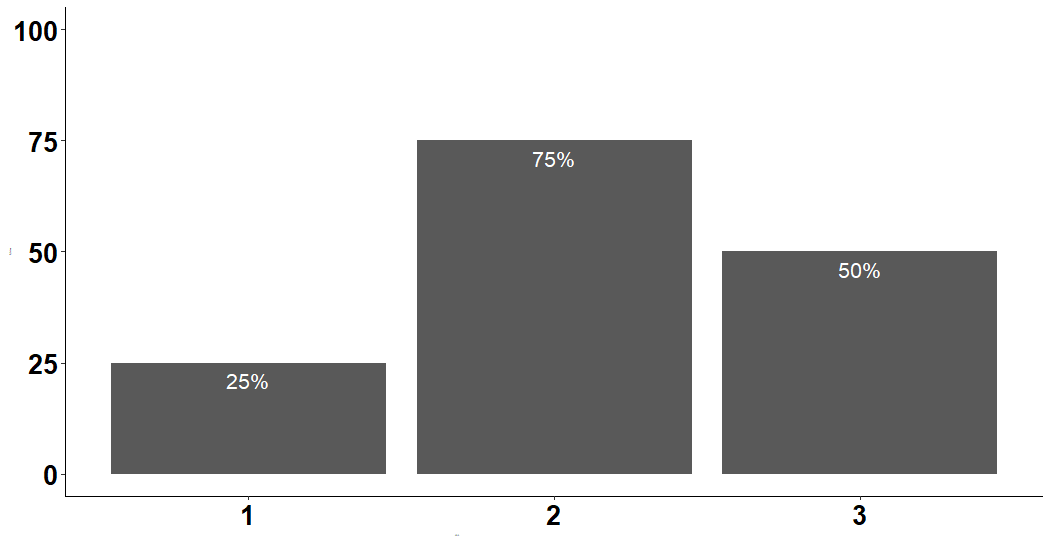
 *A. nemoralis	R. fulva*

**%**

**%**

**Taxa**

**Taxa**

1-Aphididae

2-Cecidomyiinae

3-*Diaphorina lycii*

4-*Grapholita molesta*

5*-Myzus persicae*

6-*Oenopia conglobata*

7-*Orius*

*8-Orius laevigatus*

9-*Thrips fuscipennis*

1-*Nysius graminicola*

2-*Cantharis livida*

3-Coccinellidae
